# Delineation of signaling routes that underlie differences in macrophage phenotypic states

**DOI:** 10.1101/2024.01.12.574349

**Authors:** Tiberiu Totu, Jonas Bossart, Katharina Hast, Chen Li, Markus Rottmar, Bettina Sobottka, Guocan Yu, Vanesa Ayala-Nunez, Marija Buljan

## Abstract

Macrophages represent a major immune cell type in tumor microenvironments, they exist in multiple functional states and are of a strong interest for therapeutic reprogramming. While signaling cascades defining pro-inflammatory macrophages are better characterized, pathways that drive polarization in immunosuppressive macrophages are incompletely mapped. Here, we performed an in-depth characterization of signaling events in primary human macrophages in different functional states using mass spectrometry-based proteomic and phosphoproteomic profiling. Analysis of direct and indirect footprints of kinase activities has suggested PAK2 and PKCα kinases as important regulators of *in vitro* immunosuppressive macrophages (IL-4/IL-13 or IL-10 stimulated). Network integration of these data with the corresesponding transcriptome profiles has further highlighted FOS and NCOR2 as central transcription regulators in immunosuppressive states. Furthermore, we retrieved single cell sequencing datasets for tumors from cancer patients and found that the unbiased signatures identified here through proteomic analysis were able to successfully separate pro-inflammatory macrophage populations in a clinical setting and could thus be used to expand state-specific markers. This study contributes to in-depth multi-omics characterizations of macrophage phenotypic landscapes, which could be valuable for assisting future interventions that therapeutically alter immune cell compartments.

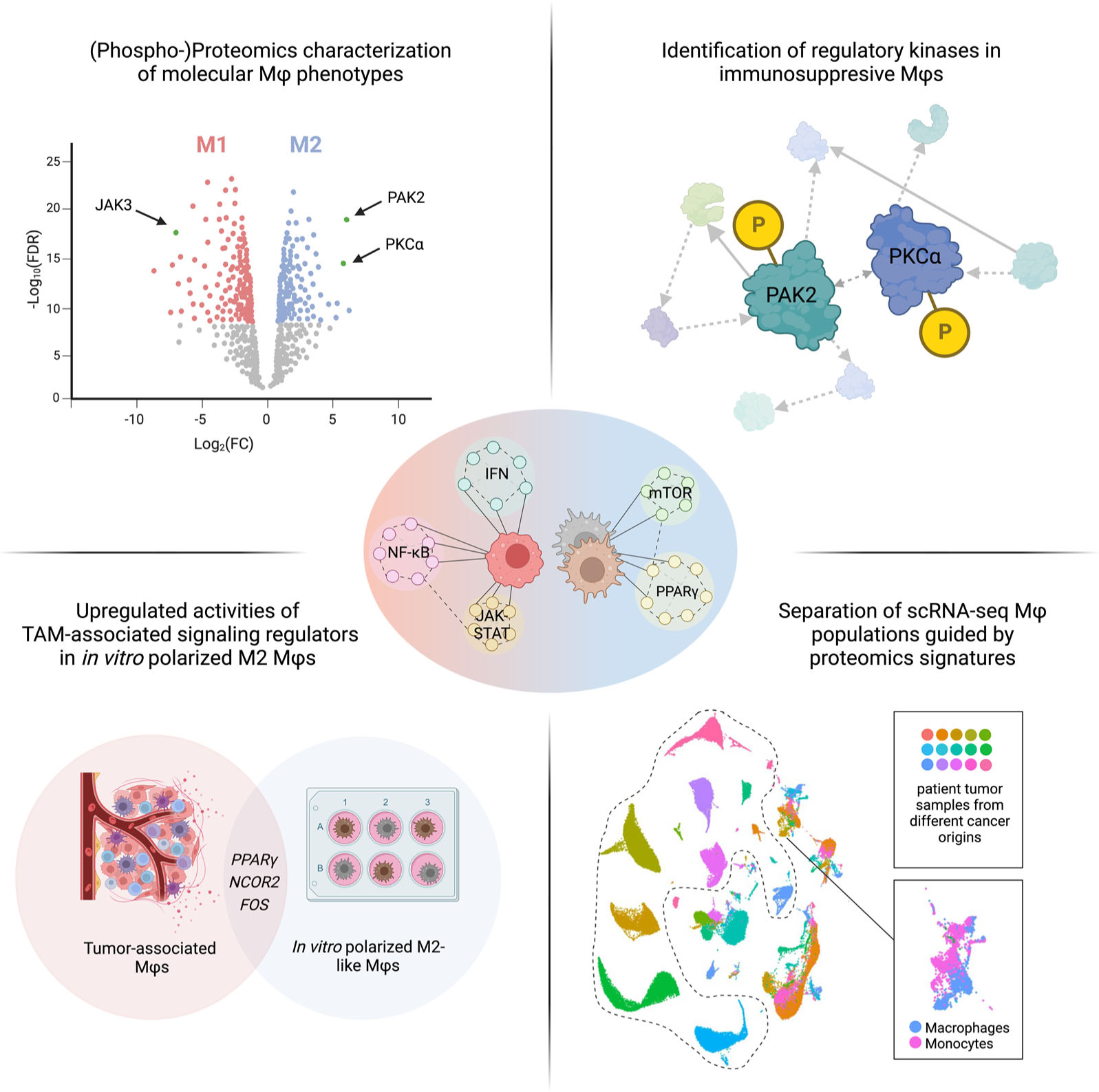

**Highlights:**

- Global proteomic characterization of primary human macrophages in different states
- Mapping of main signaling events through in-depth data analysis
- PKCα and PAK2 kinases are important regulators of immunosuppressive macrophages
- Proteomic signatures enable accurate detection of pro-inflammatory macrophages in patient tumors

## Introduction

In addition to tumor cell phenotypes, cancer progression and therapy response are strongly determined by the tumor immune microenvironment (TIME)^1^. Tumor-associated macrophages (TAMs) represent a major component of the TIMEs and mediators secreted by these cells can promote tumor viability and growth, angiogenesis and cancer invasion^2^. Macrophages are able to integrate inputs from diverse cell types in their environment and they demonstrate a high plasticity of molecular phenotypes^3^, that in effect supports the variety and flexibility of their functions. At the two ends of the broad phenotype spectrum, macrophages can have pro-inflammatory and immunosuppressive roles. *In vitro* models of these states are represented by classically activated M1 or alternatively activated M2 macrophages, respectively^4^. However, the biologically and clinically relevant macrophage spectrum entails diverse forms of macrophage states beyond the basic dichotomous M1 and M2 designations^4^. Importantly, the progression of different pathological conditions, such as cancer, inflammatory or autoimmune diseases, or the development of chronic wounds, depends on the relative fraction of macrophages in different polarization states^5^. Furthermore, due to their phenotypic plasticity, therapeutic reprogramming of macrophages is increasingly gaining attention. The development of effective and specific strategies to alter macrophage phenotypes is therefore of high interest both in basic and translational studies^6^.

In parallel with clinical and *in vivo* mouse studies, extensive research on macrophage phenotypes has been performed *in vitro*^7–9^. While *in vitro* models represent a simplified version of the dynamic and complex exposure to *in vivo* stimuli, they have proved useful for improving our understanding of macrophage biology. For instance, *in vitro* assays have been essential for determining macrophage polarization capacity of novel TIME signaling agents, such as lactate^10^, GABA^11^ or Interleukin (IL)-33^7^, or for understanding the individual signaling pathways relevant for macrophage differentiation^12^. Similarly, transcriptome data generated for macrophages, activated with a range of stimuli *in vitro*^9^, have been used for the characterization of clinically observed macrophage populations^13^. In addition, markers that relate to functionally well-characterized *in vitro* polarization states, such as CD206 after exposure to IL-4 and/or IL-13, or CD163 after exposure to both IL-4 and IL-10 stimuli are regularly used in immunohistochemistry (IHC) characterizations of clinical samples^14,15^. The ongoing studies that characterize patient TIMEs with single-cell RNA sequencing (scRNAseq) have been powerful in capturing a variety of macrophage phenotypes in a clinical setting^16,17^, but these data provide limited insight into the signaling routes that underlie the different macrophage phenotypes. In addition, mouse models have limitations as macrophage responses to pro- and anti-inflammatory stimuli are differentially regulated in humans and mice, and critical genes involved in the polarization of mouse macrophages, such as iNOS, Arg1 and TGF-β1 show different behavior in human macrophages^18^. *In vitro* assays of human macrophages are largely performed either on the differentiated THP-1 cell line or on primary blood-derived macrophages. THP-1 is a human leukemia monocytic cell line, which does not accurately represent relevant immune processes or reflect inter-individual variability in macrophage responses to stimuli^19^.

While signaling cascades that define pro-inflammatory macrophages with anti-tumorigenic roles are better characterized, regulatory pathways that drive polarization of immunosuppressive macrophages are still incompletely mapped^20,21^. Novel kinases that regulate these processes are still being discovered and are of immense interest as possible reprogramming targets, as illustrated by the recent studies of the RIP1^22^ and PI3Kγ kinases^23,24^. Previous investigations of signaling pathways that are of relevance in TAMs relied on catalogues generated by independent characterizations of *in vitro* polarized macrophages where antibody assays against a small set of signaling phosphoproteins were used^7^. Such targeted assays detect only a small fraction of events found with large-scale unbiased mass spectrometry (MS)-based phosphoproteomics approaches^25,26^. MS-based proteomics and phosphoproteomics assays have so far been mostly performed on murine macrophages or on macrophages generated from a THP-1 cell line^27–30^. Even though proteomes of primary human macrophages are also available, detailed analysis of relevant signaling events has not yet been reported^31–33^. Primary macrophages have been characterized by several transcriptomic studies^34–42^. However, transcriptomics data cannot pinpoint the exact components of cellular signaling cascades and directionality of signal flow, whereas phosphoproteomics allows the simultaneous and unbiased assessment of hundreds of kinases, either by direct measurement of their phosphorylation status or by the footprints of their activities.

Crucial signaling mediators for M1 *in vitro* (and M1-like *in vivo*) polarized pro-inflammatory macrophages are well characterized and it has been consistently reported that the main signaling routes after exposure to Interferon γ (IFN-γ) and Lipopolysaccharide (LPS) stimuli lead to the activation of p38, JAK1 and JAK2 kinases and higher expression of genes regulated by the STAT1, STAT5, IRF3, IRF5, IRF8 and NF-κB transcription factors (TFs)^43^. Immunosuppressive macrophage states polarized *in vitro* are described as M2a and M2c phenotypes when they are generated after exposure to IL-4 and IL-13, or to IL-10, respectively. M2b state requires stimuli with immune complexes and IL-1β, and is not characterized as immunosuppressive^44^. M2a macrophages are characterized by high activity of IRF4 and STAT6 TFs as well as with increased activity of retinoic acid pathway and signaling from AKT and MAPK kinases^27,45^. The M2c state is less well studied than the M2a, but is known to have increased activity of the STAT3 TF and AMPK kinase, as well as an upregulation of SOCS3, CXCL13 and several metalloproteases^46^. Immunosuppressive macrophages have roles in tissue healing, activation of regulatory T-cells, matrix remodeling and angiogenesis^34,45^.

Here, we performed a global characterization of signaling events in primary human M1, M2a and M2c macrophages using MS-based proteomic and phosphoproteomic profiling. We compared signaling activities in pro-inflammatory phenotypes stimulated with LPS and IFN-γ (known as M1) and immunosuppressive phenotypes stimulated either with IL-4 and IL-13 (M2a) or with IL-10 (M2c). Analysis of direct and inferred footprints of kinase activities indicated a high activity of RIPK2, SRC and JAK2/3 kinases in pro-inflammatory phenotypes and it suggested PKCα, PAK2, LRRK2 and MAST kinases as likely regulators of immunosuppressive macrophages. By integrating these data together with publicly available transcriptomics datasets on macrophages exposed to the same stimuli and by using network modularization on a comprehensive framework of known physical and genetic interactions among these proteins, we extracted interaction neighborhoods specific to each phenotype of interest. This showed that protein modules, which associate with the main signaling routes of mTOR and MAPK pathways, had the most significant changes across the studied phenotypic states. Finally, we used proteome signatures identified here to classify macrophages in tumor samples from cancer patients, which were characterized with scRNAseq, and found that the proteomic markers were able to successfully distinguish pro-inflammatory macrophage populations in a clinically relevant context. Overall, in-depth multi-omics characterization of macrophage phenotypic landscapes could be valuable for supporting the rational design of interventions that aim to therapeutically alter immune cell compartments.

## Results

### Macrophages polarized to different states express specific protein markers

In order to systematically investigate signaling pathway differences between differentially polarized human macrophages, we obtained buffy coats from four blood donors and isolated the CD14^+^ monocytes (Figure 1A, see Methods). Following, polarization to the M1 state was achieved using LPS and IFN-γ as stimuli (after differentiation with GM-CSF), while M2 states were generated either with IL-4 and IL-13 (corresponding to the *in vitro* M2a annotation) or with IL-10 (corresponding to the *in vitro* M2c annotation, both after differentiation with M-CSF, see Methods)^8,47^ (Figure S1A). In order to validate polarization to M1 and M2 phenotypes, we used flow cytometry and assessed the expression of cell surface proteins which are commonly used as markers for the studied states^48–51^: CD86 for M1, CD206 for M2a and CD163 for M2c. This showed that cell populations exposed to one of the three stimuli conditions indeed had a higher expression of the markers for the corresponding cell states (Figure 1B, Figure S1B). M1 cells had the highest fraction of CD86 positive cells (79 and 85% in the two tested donors), M2a cells had the highest fraction of cells positive for the CD206 marker (56% in both tested donors) and M2c cells had the highest fraction of cells positive for the CD163 marker (74 and 91% in the two tested donors). This indicated that the achieved polarization states represented those previously described in the literature.

**Figure 1.**
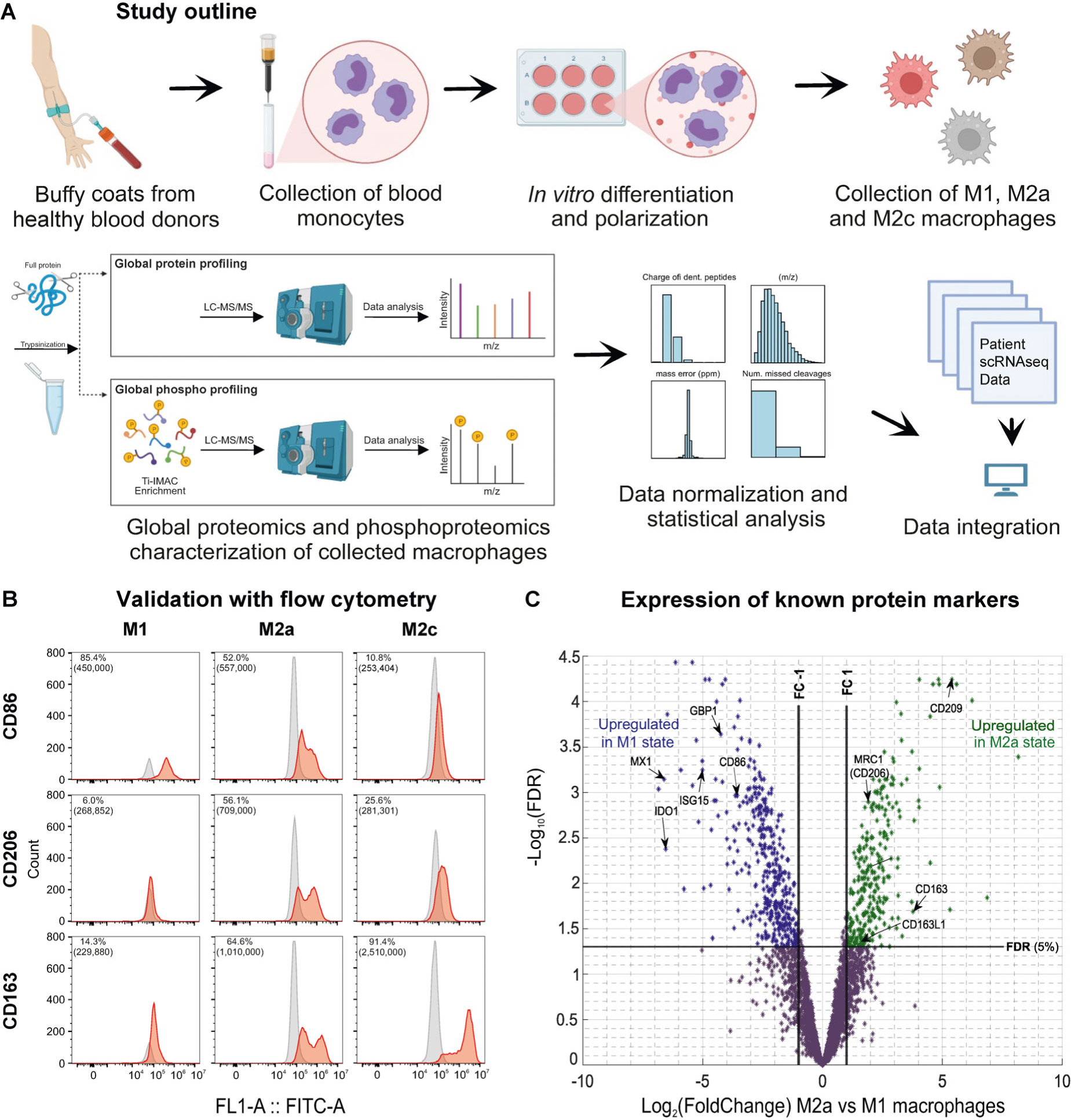
Generation and validation of *in vitro* polarized M1, M2a and M2c phenotypic states for primary human macrophages. (A) Macrophage polarization and (phospho-)proteomics analysis workflow for comparative analysis of primary human M1, M2a and M2c macrophages. (B) Flow cytometry profiles for the polarization markers CD86, CD206 and CD163 of *in vitro* polarized macrophage phenotypes are shown in red and negative or isotype controls are shown in grey. Percentage of positive cells and the median fluorescence intensities are indicated on each plot. The latter is shown in brackets. Y-axes are truncated. (C) Volcano plot illustrating protein expression in terms of Log_2_FC values on the X axis against −Log_10_(FDR) values on the Y axis in a comparison of the M1 and M2a phenotypic states. Differentially expressed proteins are colored blue and green, depending on the directionality of the expression change.

We performed global proteomic and phoshoproteomic characterization of the stimulated cells (see Methods). For the latter, phosphopeptides were enriched with Ti-IMAC microparticles. Of note, protein phosphorylation can have either an activating or an inhibitory effect on the protein. We used the MaxQuant software tool to match peptides to proteins and estimate quantities of the measured analytes (see Methods)^52,53^. The label-free proteome and phosphoproteome quantification resulted in the identification of a total of 5,342 proteins and 5,905 phosphopeptides, which mapped to 2,313 phosphoproteins. We first compared protein levels in the M1 cells to those in the M2a and M2c cells. In total, we found that 675 and 806 proteins had significantly different expression levels between M1 and M2a, and between M1 and M2c macrophages, respectively (two-tail moderated *t*-test, false discovery rate (FDR) < 0.05 and Log2 FoldChange (FC) > 1, Figure 1C and Figure S1C, Table S1). Of these, 282 and 299 proteins were expressed at significantly higher levels in M2a and M2c, respectively, when compared to M1 macrophages.

Next, we used Gene Ontology together with KEGG and Reactome pathway annotations in order to identify the major functional roles of the differentially regulated proteins (see Methods and Table S2). Proteins that were significantly upregulated in the M1 phenotype were strongly enriched in the members of interferon and cytokine signaling pathways (FDR < 6.40 × 10^−9^ for the *Interferon Signaling* and FDR < 1.62×10^−4^ for the *Cytokine Signaling in Immune system* Reactome pathways, Figure S1D), while proteins upregulated in the M2 phenotypes were enriched in metabolic roles (FDR < 2.21×10^−3^ for the *Amino sugar and nucleotide sugar metabolism* KEGG pathway and FDR < 4.71×10^−2^ for the *Metabolism of carbohydrates* Reactome pathway). Among the proteins that were highly upregulated in the M1 macrophages compared to both M2a and M2c macrophages (Table S1) were CD86, which is used as a surface marker for the M1 state (FDR < 1.09×10^−3^ and Log_2_FC > 3.38), as well as a number of well-studied inflammatory proteins, such as GBP1, IDO1, ISG15, and MX1^54,55^ (GBP1: FDR < 2.31×10^−4^ and Log_2_FC > 4.12; IDO1: FDR < 4.21×10^−3^ and Log_2_FC > 6.54; ISG15: FDR < 5.66×10^−4^ and Log_2_FC > 5.01; MX1: FDR < 7.18×10^−4^ and Log_2_FC > 5.70). The respective proteins are marked in Figure 1C and Figure S1C. Analogously, IL-4 and IL-13 treated macrophages had highly upregulated levels of several surface proteins that were used in previous studies as markers for immunosuppressive M2a macrophages. These included CD209, CD206/MRC1 and CD163 (CD209: FDR < 5.74×10^−5^ and Log_2_FC > 5.39; MRC1: FDR < 1.27×10^−3^ and Log_2_FC > 1.90; CD163: FDR < 2.05×10^−2^ and Log_2_FC > 3.77)^28,56,57^. Similarly, M2c macrophages had a significantly upregulated expression of surface markers CD209 and CD163 (CD209: FDR < 4.93×10^−4^ and Log_2_FC > 3.36; CD163: FDR < 2.63×10^−3^ and Log_2_FC > 5.47). Overall, this showed that the *in vitro* polarized M1 and M2 macrophages studied here could be clearly distinguished from each other through a differential expression of proteins associated either with pro-inflammatory or with immunosuppressive phenotypes.

### Several regulatory proteins, which are expressed in TAMs and known to promote tumor growth, have elevated phosphorylation levels in M2 macrophages

Analogously to the protein level analysis, we assessed which phosphopeptides showed significant differences in their quantitative levels between the M1 and M2 states (Figure 2A and Figure 2B). In order to be able to interpret the observed changes as a result of the higher activity of the upstream kinase(s), we considered only phosphoproteins for which changes in quantitative levels of at least one of their phosphopeptides could not be explained by the expression level changes of the corresponding protein (see Methods). In this way, the comparison of M1 and M2a phenotypes highlighted 1,986 phosphopeptides (out of 5,880 measured ones) with significantly different quantitative levels between the two states (FDR < 0.05 and Log_2_FC > 1, two-tail moderated *t*-test). With the same FDR and Log_2_FC thresholds, in total 2,780 phosphopeptides (out of 5,153 measured ones) were found to be differentially regulated between the M1 and M2c phenotypes. We compared proteins with significant phosphorylation changes to the background of all measured phosphoproteins and assessed their functional enrichment in the components of the KEGG and Reactome signaling pathways (see Methods and Figure S2A). This showed that proteins with higher phosphorylation levels in M1 were, among others, significantly enriched in the members of the Interferon gamma signaling pathway (FDR < 1.49×10^−2^), whereas those with higher levels in M2a were enriched in the components of the mTOR signaling pathway (FDR < 1.12×10^−3^). Proteins with higher phosphorylation in M2c were enriched in Fcγ mediated phagocytosis (FDR < 4.62×10^−2^). M2c macrophages have been previously reported to have a high phagocytic capacity^58,59^, while the mTOR pathway, which has a crucial role in the regulation of cellular metabolism and proliferation, is considered to be at the crossroad between M1 and M2a polarization^60^.

**Figure 2.**
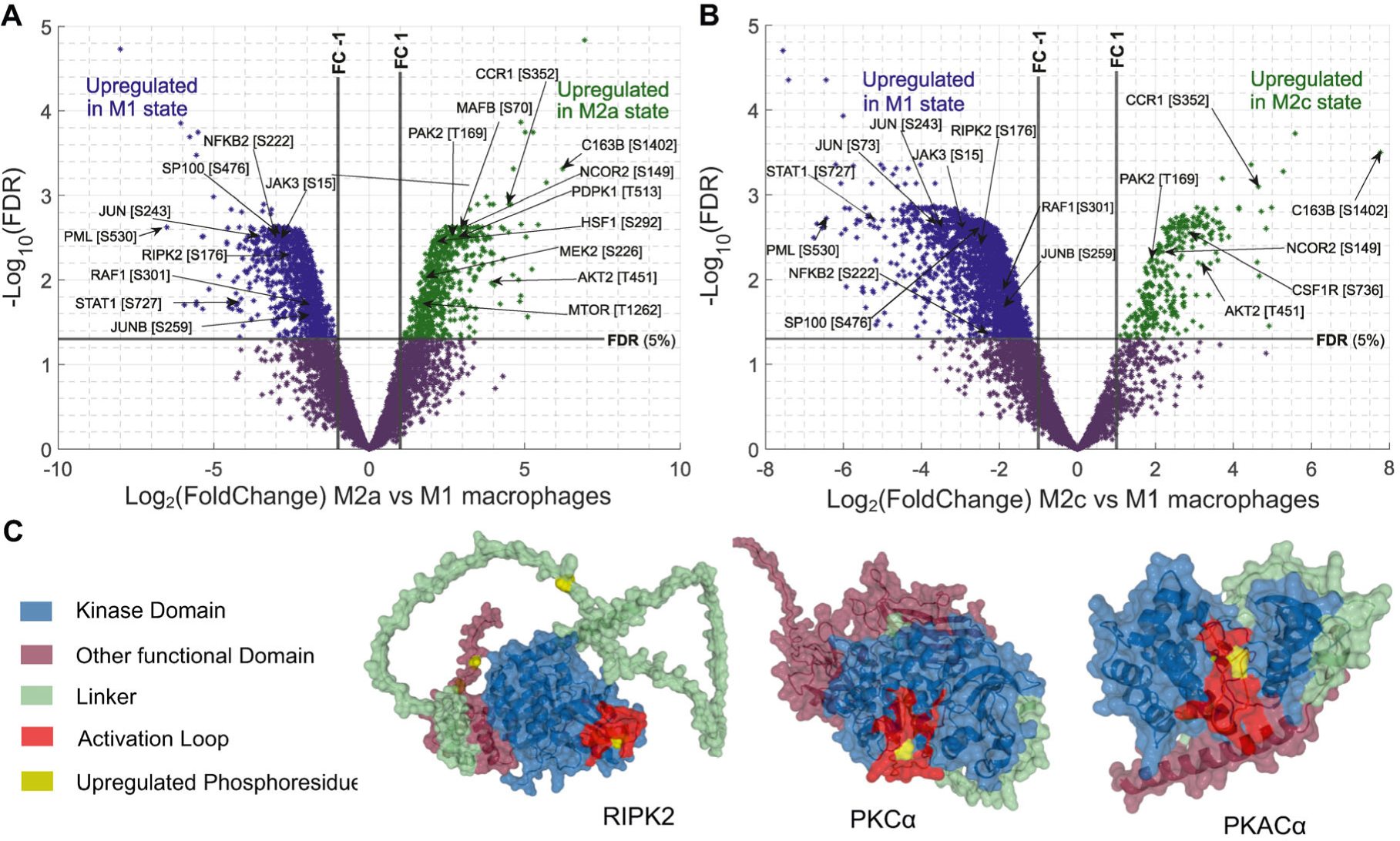
Phosphoproteomics analysis highlights phenotype-state specific upregulation of specific kinases and other signaling proteins. (A) Volcano plot comparing peptides with different phosphorylation levels between *in vitro* polarized M1 and M2a macrophage phenotypes. A Log_2_FC of 1 and an FDR value of 5% in a moderated *t*-test were used as thresholds for significant hits. (B) Volcano plot comparing peptides with different phosphorylation levels between *in vitro* polarized M1 and M2c macrophage phenotypes. (C) 3D protein structure of example kinases phosphorylated in their activation loops. RIPK2 was found to have increased phosphorylation level in the M1 inflammatory macrophages and PKCα in M2a immunosuppressive macrophages. In both proteins, phosphosite maps within the active loop, while PKACα is a highlight for M2c. The phosphoresidues measured and found upregulated in the studied phenotypes are marked in yellow while different protein functional regions are indicated according to the legend. Structures were rendered in PyMOL version 2.5.2.

Proteins with the most significant differences in phosphopeptide levels between M1 and M2 states (Figure 2A and Figure 2B, Table S1) included the tumor suppressor PML, which had 12 phosphoresidues with higher phosphorylation levels in the M1 state compared to both M2a and M2c macrophages (all 12 sites with an FDR < 4.85×10^−2^ and Log_2_FC > 1.99). PML has multiple roles in the formation of PML-nuclear bodies and it is linked to the IFN-γ signaling pathway^61^. In addition, tumor suppressor SP100, which together with PML is a major constituent of the PML bodies, also had eight phosphosites with higher levels in M1 macrophages (FDR < 3.81×10^−2^ and Log_2_FC > 1.67)^62^. There were 191 phosphosites upregulated in both M2a and M2c macrophages when compared to M1 (Figure S2B). These included phosphoresidues of the CCR1 chemokine receptor, which was previously shown to promote M2 macrophage polarization^63^ (FDR < 7.07×10^−3^ and Log_2_FC > 2.13). In addition, the NCOR2 TF, which is able to suppress inflammation, showed higher phosphorylation in the M2 states^64,65^ (S149 and S152: FDR < 4.78×10^−3^ and Log_2_FC > 2.12). In the M2a state, MAFB and HSF1 TFs, whose elevated expression in TAMs associates with more aggressive tumor growth^66,67^, had significantly upregulated phosphosites (MAFB S70: FDR < 2.39×10^−3^ and Log_2_FC > 2.71; HSF1 S292: FDR < 3.45×10^−3^ and Log_2_FC > 2.27).

A central element of cellular information flow is through phosphorylation of protein kinases by other kinases. Therefore, we specifically investigated differences in kinase phosphorylation levels across the studied states. In total, 71 protein kinases had significantly different quantitative levels of one or more phosphosites in the comparison between M1 and M2 phosphoproteomes. We investigated if these kinases were enriched in specific KEGG pathways and found that 21 of the 71 significant kinases belonged to the MAPK signaling pathway (this presented an enrichment when compared to all measured kinases; FDR < 4.90×10^−2^, modified Fisher’s test), 14 kinases belonged to the Chemokine signaling pathway (FDR < 1.01×10^−2^) and 13 to the mTOR signaling pathway (FDR < 9.20×10^−3^). Among the kinases with significantly higher phosphosite levels in the M1 state compared to either of the M2 states were JAK2 (S518: FDR < 4.97×10^−3^ and Log_2_FC > 2.70) and JAK3 kinases (S17 and S15: max FDR < 3.43×10^−2^ and Log_2_FC > 2.12), which are known to form the main axis of the JAK-STAT pro-inflammatory pathways. Specifically, the phosphorylation of S518 in JAK2 was previously shown to play a role in the kinase activation^68^. In addition, SRC and RAF1 kinases had phosphoresidues with a higher phosphorylation level in M1 (SRC S43: FDR < 4.24×10^−2^ and Log_2_FC > 1.53; RAF1 S301: FDR < 1.87×10^−2^ and log_2_FC > 1.89). SRC has been reported to play a role in the production of inflammatory cytokines and mediators in macrophages^69^ and RAF1 regulates the MAPK pathway, which is important in the M1 state^70,71^. Kinases with higher phosphorylation levels in the M2a state included MAST2 and MAST3 kinases, where MAST2 is known to be able to block the NF-κB activation^72,73^ (MAST2 S1364: FDR < 1.36×10^−2^ and Log_2_FC > 2.0; MAST3 had four upregulated residues, all with FDR < 3.89×10^−3^ and Log_2_FC > 2.56). In addition, several other kinases had significantly higher phosphorylation levels in the M2a state: PAK2 kinase, which was previously reported to regulate the development of myeloid-derived suppressor cells in mice^74^, WNK1 kinase, which is able to suppress inflammatory cytokine production^75^, LRRK2 kinase, which regulates different inflammatory responses in the body^76^, and PDPK1 kinase, which is known to regulate the mTOR pathway and promote M2-like polarization in mice^77^ (PAK2 T169 and S58: FDR < 5.0×10^−3^ and Log_2_FC > 2.57; WNK1 S2027: FDR < 2.39×10^−3^ and Log_2_FC > 2.42; LRRK2 S973: FDR < 1.33×10^−2^ and Log_2_FC > 1.75; PDPK1 T513: FDR < 3.28×10^−3^, Log_2_FC > 2.67). In the M2c state, we observed a higher phosphorylation of the FES kinase (T421: FDR < 3.32×10^−3^ and Log_2_FC > 2.39), a widely expressed kinase, which is able to downregulate the immune response during inflammation^78,79^.

Phosphorylation within the kinase activation loop or an analogous regulatory segment is often sufficient for the kinase to switch to its active state. Several of the phosphosites with significant differences in their phosphorylation levels across the macrophage states mapped within sequence segments annotated as regulatory regions (Figure 2C and Figure S2C). For instance, the S176 phosphoresidue within the RIPK2 kinase had a significantly higher phosphorylation level in the M1 state when compared to both M2 states (FDR < 4.72×10^−3^ and Log_2_FC > 2.51). The phosphosite maps within the RIPK2 activation segment, and the S176 residue itself is annotated as an auto-phosphorylation site essential for the RIPK2 catalytic activity^80,81^. RIPK2 plays an important role in the activation of pro-inflammatory pathways, including NOD signaling and NF-κB pathways^82^. Furthermore, the activation residue of the PKCα kinase (a.k.a PRKCA)^83^, T497, had higher phosphorylation levels in both M2 states when compared to M1 (max FDR < 4.66×10^−2^ and Log_2_FC > 1.28). The PKCα kinase is known to play a role in anti-inflammatory processes and it may negatively regulate the NF-κB induced genes^84^. Finally, in the M2c state, the PKACα (a.k.a PRKACA) T198 phosphosite was measured at a significantly higher level than in the M1 state (FDR < 3.86×10^−3^ and Log_2_FC > 2.72). The residue maps within the kinase activation loop and its phosphorylation is associated with the increase in the kinase’s catalytic activity^85,86^. PKACα was reported to be able to induce a pro-tumoral immunosuppressive macrophage phenotype^87^. Jointly, these results show that phosphoproteome characterization of macrophage functional states is able to recognize known molecular mechanisms that underlie their different functional roles and highlight a number of novel instances, which link to signaling pathways that can drive pro-inflammatory and immunosuppressive cell phenotypes.

### Analysis of kinase activity footprints implies possible regulatory roles for LRRK2, PKCα and PAK2 kinases in immunosuppressive macrophage states

The main advantage of quantitative phosphoproteomics, compared to transcriptomics or proteomics, is that it provides closer insights into the active states of proteins. However, phosphoproteomics is still hampered by a high fraction of missing values and the absence of a certain phosphoprotein does not necessarily mean that the protein is not present in the phosphorylated form^88^. In order to infer the increased activity of upstream protein kinases, we further studied sequence motifs surrounding all significantly upregulated phosphosites and considered also upstream kinases that were themselves not measured. For this, we made use of The Kinase Library, a recently published analytical tool based on a systematic screen of synthetic peptide libraries^89^.

This analysis indicated pro-inflammatory JNK1, JNK2 and JNK3 kinases (FDR < 2.38×10^−2^), together with p38 mitogen-activated kinases (FDR < 9.72×10^−2^), as major regulators of phosphoproteome changes in the M1 macrophages (Figure 3A and Table S3). Thus, the Kinase Library analysis, even though based solely on phosphopeptide sequences, correctly predicted known major signal transduction routes in the M1 state^90^. Analysis of phosphoresidues upregulated in the M2a state suggested a high activity of IRAK1 (FDR < 1.07×10^−4^) and IRAK4 (FDR < 2.42×10^−2^) kinases (Figure 3B and Table S3). Together with the TRAF6 protein, these kinases can form a complex, that activates pro-inflammatory JNK kinases and the NF-κB TF^91^. However, when the IRAK1/4 complex binds other partners, such as IRAK-M, it can act as a negative regulator of inflammation and phosphorylate a different set of downstream substrates^91,92^, which could explain its increased activity in the M2a state. This analysis also suggested a higher activity of the LRRK2 (FDR < 2.27×10^−2^) and GAK (FDR < 1.63×10^−2^) kinases in M2a macrophages (Figure 3B). LRRK2 has been linked to pathways that regulate inflammation^76^ and the less well-studied GAK kinase is one of its few confident interaction partners^93^. In M2c macrophages, the Kinase Library analysis suggested upregulation of the GAK (FDR < 6.75×10^−2^) and CAMKK2 (FDR < 4.75×10^−3^) kinases (Figure 3C). High expression levels of CAMKK2 were previously reported in TAMs and activation of this kinase was shown to support tumor growth^94^.

**Figure 3.**
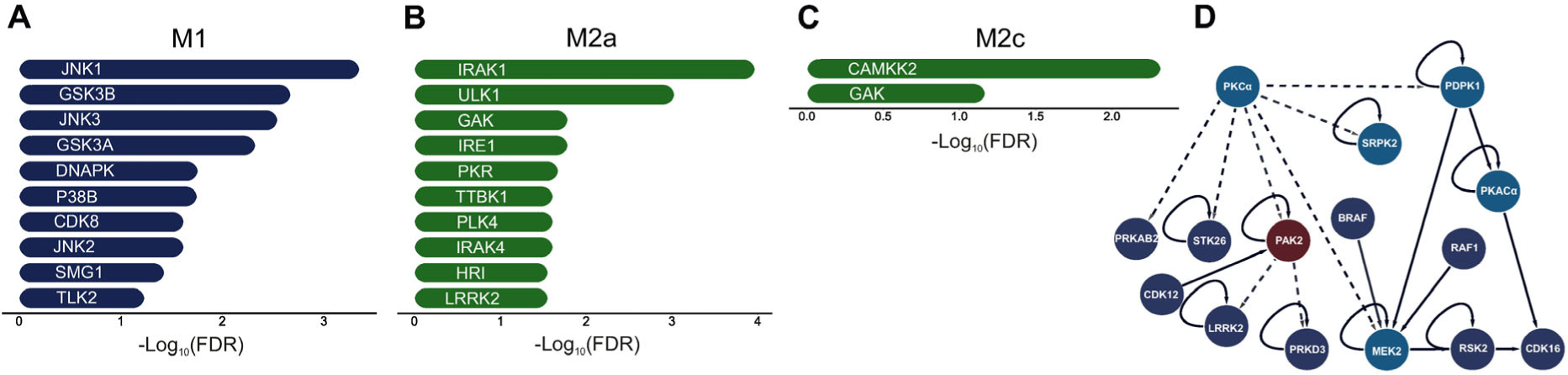
Inferred upstream kinases responsible for the phosphorylation of upregulated phosphopeptides highlight both known and novel kinases and signaling routes. (A-C) Kinases with the most significant predictions for the different activity levels between the M1 and M2 phenotypes are shown. The Kinase Library tool was used for this. There, *p*-values are calculated with the one sided Fisher’s exact test and corrected for multiple testing with the Benjamini-Hochberg method. (A) Kinases inferred as significantly upregulated in the M1 state when compared to M2a. All measured phosphosites were considered, together with their Log_2_FC between the states and FDR values calculated with the moderated *t*-test and corrected for multiple testing. (B) Kinases inferred as significantly upregulated in the M2a state when compared to M1. (C) Kinases inferred as significantly upregulated in the M2c state when compared to M1. (D) Kinases with increased phosphorylation levels in the activation loop (light blue) in the M2a macrophages are shown together with their first-degree kinase neighbors (dark blue). In addition, first-degree kinase neighbors of the PAK2 kinase are included due to NetPhorest predictions for its increased activity in M2 macrophages. All kinases shown here have at least one upregulated phosphopeptide in the M2a state (with respect to the M1 phenotype). All interactions indicate kinase-substrate relationships. Solid lines show those obtained from the curated knowledge databases (such as PhosphositePlus) and dotted lines show those from the NetPhorest prediction tool.

In addition, we conducted further analyses for complementary predictions of upregulated kinases: (i) we applied the NetPhorest tool^95^, which also investigates phosphosite sequence motifs and (ii) we used the Kinase Enrichment Analysis version 3 (KEA3) method for the analysis of phosphoproteome datasets, which further considers known protein interactions and co-expression trends^96^. Analysis of known kinase-substrate relationships from the curated annotations collected in the PhosphoSitePlus and other databases^97^ did not yield clear trends (Table S3). Nevertheless, the curated annotations, alongside the predicted ones, facilitated the construction of kinase-kinase signaling networks that describe the signaling flow within the studied phenotypes as was measured in this study (Figure 3D and Figure S3A-C). The NetPhorest analysis recapitulated a high activity of kinases from the p38 family in the M1 state (FDR < 3.39×10^−2^, Figure S3D). Furthermore, it highlighted the upregulation of kinases from the PKC group in the two M2 states (FDR < 5.25×10^−3^, Figure S3E and Figure S3F). The PKC group contains the above discussed PKCα kinase, which we observed to have the activation loop phosphorylated in the M2 states. Finally, the KEA3 predictions (Table S3) included JAK and MAPK kinases as top hits for the M1 phenotype, as well as PKACα and PAK2 kinases as hits for both the M2a and M2c phenotypes. As discussed above, the PAK2 kinase itself had two phosphosites significantly upregulated in the M2a macrophages, while PKACα was found to be phosphorylated in the activation loop. Jointly, analysis of kinase signal propagation in primary macrophages *in vitro* has further highlighted known routes relevant for the establishment of polarized states and suggested several novel kinases that could play a decisive role in the different immunosuppressive macrophage phenotypes.

### Network integration of macrophage omics data points towards central pro-inflammatory and immunomodulatory protein modules

Several previous studies have performed transcriptomic analysis of primary human macrophages polarized with the same stimuli as here^34–39,98^. We retrieved the published datasets from the six studies and re-analyzed the data in order to systematically assess cell state-specific differences in molecular activity. All of the studies included M1 and M2a phenotypic states with two of them additionally including M2c macrophages. For the M1 and M2a comparison, we included in the final list of significantly differentially expressed genes those that were found as such in at least three studies. In this way, we identified 1,252 high-confidence polarization state-specific genes (FDR < 0.05, abundance ratio > 4, see Methods and Table S4). Differentially expressed genes between M1 and M2c states were were selected with the same threshold criteria, but were defined as a less stringent union of the two available studies (2,911 genes in total). We performed functional enrichment analysis and found that genes with higher expression levels in the M1 phenotype were enriched in the KEGG and Reactome pathways associated with Interferon, TNF and NF-κB signaling (with FDR < 1.51×10^−26^, FDR < 6.62×10^−10^ and FDR < 8.75×10^−8^, respectively). Genes with a higher expression in the M2a and M2c macrophages were enriched in the PPAR signaling pathway (FDR < 1.71×10^−2^), which was previously linked to the M2 phenotypes^99,100^.

Next, we searched for the likely upstream regulators of the high confidence differentially expressed genes by using annotations on known targets of human TFs available in the TRRUST database^101,102^. Genes upregulated in the M1 state indicated that the major TFs with M1-increased activity were RELA, NFKB1 and STAT1, all well-known regulators of this state (FDR < 8.49×10^−6^, Hypergeometric test, Table S4). Upstream TFs in the M2a state were detected with a lower significance (FDR < 0.15), reflecting both less strong gene upregulation and fewer annotations on these genes. However, the top hits for M2a were STAT6, KLF2 and ETS1 TFs. STAT6 is a known marker of M2a macrophages, while KLF2 is a negative regulator of pro-inflammatory genes^103^. In addition, we used the same datasets to identify genes with differential transcript usage between the M1 and M2a states and found 697 genes with differential alternative splicing (Table S4, see Methods). When compared to all other genes with measured transcripts, these genes had more often significant changes in phosphopeptide levels (FDR < 8.8×10^−4^, Fisher’s exact test).

Following, we used the obtained catalogue of differentially expressed genes, proteins and phosphoproteins as well as genes with differential transcript usage in order to assess polarization state-specific cellular networks. To build the networks, we integrated confident interactions collected from the STRING, BioGRID and IntACT databases (see Methods). We retained only interaction pairs where both of the proteins were detected as differentially expressed in the same state with at least one type of analysis. Networks built this way allowed us to i) identify central network elements for each state and ii) identify network modules of highly connected proteins that likely share similar functional roles. For the former, we used the current flow betweenness centrality metric, which assesses the shortest paths that connect network elements through each respective protein, and additionally includes contributions from all possible paths by accounting for the information flow through random walks^104^. When considering networks with hits upregulated in individual states, 10% of the proteins with the highest centrality measure contained 87, 48 and 67 proteins in M1, M2a and M2c states, respectively (Table S5). Central proteins in the M1 state, which were found consistently upregulated in different omics analyses, included a number of signaling regulators that are known to play a crucial role in M1 polarization, such as STAT1, STAT3, RELA, JUN, NFKB2 and NCOR1 TFs as well as SRC, JAK2, JAK3, RIPK2 and MAPK11 kinases and the above-mentioned PML protein. Network hubs in the M2a state included transcription regulators FOS and PPARɣ. In addition, the transcription regulator NCOR2 was among the top 25% of the M2a most significant hubs. FOS was reported to be able to suppress inflammation^105^ and the PPAR pathway is one of the hallmarks of M2a macrophages^27^. FOS TF was also identified as a hub in M2c macrophages. Furthermore, in both M2a and M2c states, the CSF1R receptor was identified as one of the central regulators. CSF1R is able to direct monocyte migration to tumors and promote M2-like polarization *in vivo*^106^. Several antibodies and inhibitors that block its activation are considered in clinical trials that aim to revert tumor immunosuppression^107^. In addition, this analysis highlighted mTOR, PKCα, PAK2 and LRRK2 kinases as central regulators in M2a macrophages. Of note, the PKCα kinase was a protein with the highest centrality score in the M2a network. Jointly, these results suggest that integrated network analysis is able to identify candidates that warrant further investigation for their regulatory roles in distinct macrophage states.

In order to perform network decomposition, we applied a modularity optimization algorithm within the MONET toolbox^108^, which is able to identify network communities at different resolutions. We used the same networks as above with hits identified as significant either through previous transcriptomic studies or though proteomic and phosphoproteomic datasets generated here. For the M1, M2a and M2c states, we were able to distinguish 14, 17 and 3 confident modules, respectively, each with 10 or more members (Table S5). Modules for the better studied M1 macrophages distinguished protein communities with dominant roles in IFN-γ (first module), and in NF-κB, toll-like receptor and JAK-STAT signaling (second and third modules). The largest module detected in the M2a state (first module) contained mTOR and MAPK pathway components and included PAK2, MEK2 and PDPK1 kinases (Table S5). As expected, there was a high interconnectivity among the module components. This is illustrated in Figure 4, which shows links between transcription factors with a high centrality in the M1 and M2a states together with their interaction partners that had the highest current flow betweenness centrality scores. Overall, cellular maps constructed with significant proteins from multi-omics analyses were able to effectively summarize our knowledge of proteins with central roles in macrophage polarization, add new members to the better studied functional modules, and highlight protein communities that could have additional roles in promoting distinct macrophage states.

**Figure 4.**
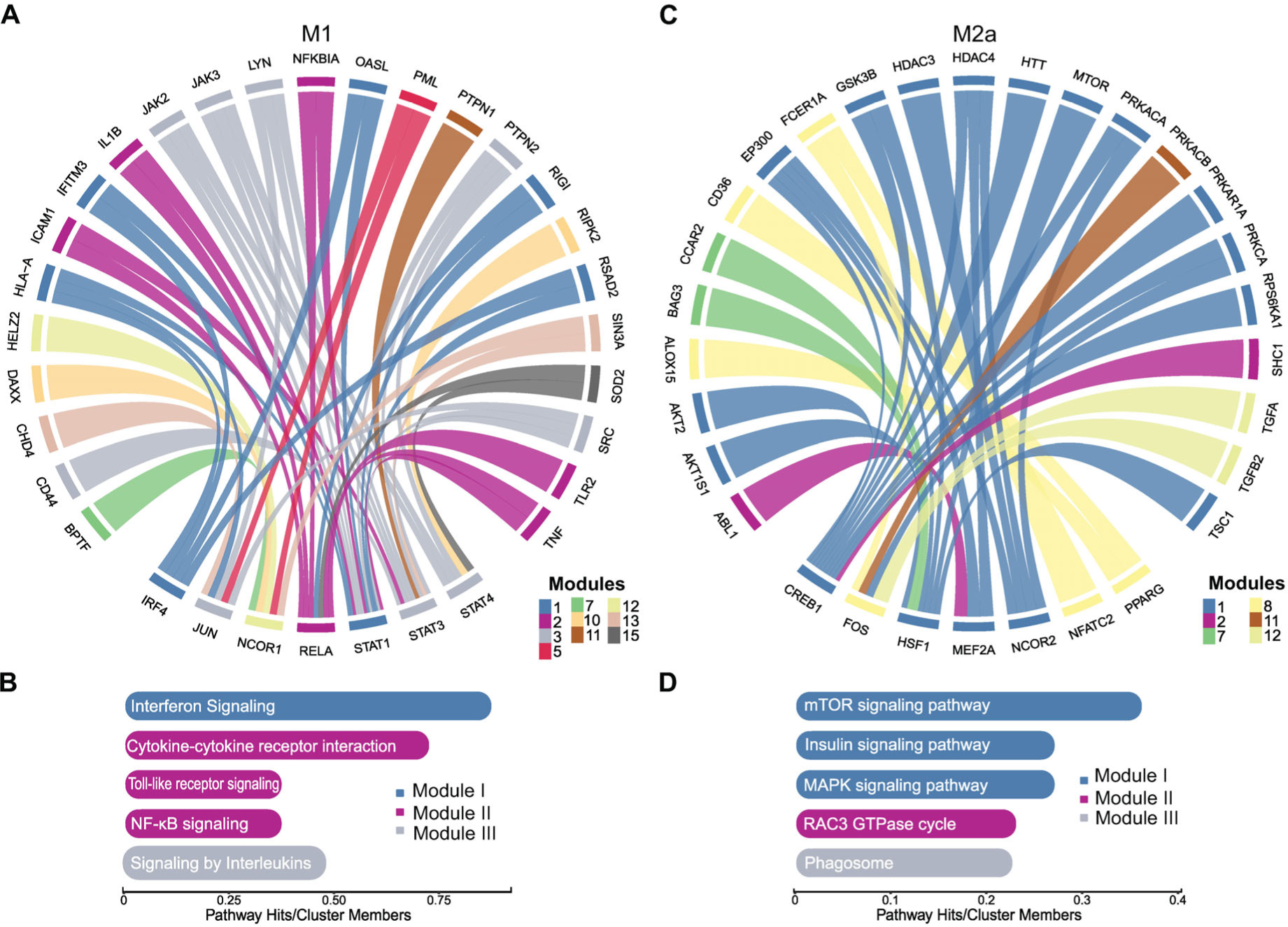
Central elements in the integrative network built from the genes and proteins found upregulated in the proteomics, phosphoproteomics and transcriptomics analysis point to state-specific signaling pathways. (A) Circular diagram illustrates TFs and their interaction partners which were defined as most central nodes based on the current-flow betweenness centrality score. On the lower side of the plot are the TFs with their interaction partners shown above. For readability, 7 most central TFs and maximum 25 of their partners with the highest centrality scores are shown. Diagram coloring is based on the module these proteins were assigned to after the network decomposition. In this plot, only entries found upregulated in the M1 state (when compared to M2a) in either of the omics analysis are shown. (B) Pathway enrichments of proteins assigned to different modules after network decomposition analysis for the M1 phenotype. All shown terms were found significant with an FDR < 0.05 (hypergeometric test with Benjamini-Hochberg correction for multiple testing, when compared to all proteins in the network). A fraction of proteins in the module with the respective annotation is shown. (C) Analogous as in (A). Here, only entries with a high centrality score that were found upregulated in the M2a state (when compared to M1) in either of the omics analysis are shown. Again, 7 most central TFs and maximum 25 of their partners with the highest centrality scores are shown. (D) Pathway enrichments of proteins assigned to different modules after network decomposition analysis for the M2a phenotype. Data is presented analogous as in (B).

### Proteomic markers successfully distinguish pro-inflammatory macrophages in a clinical context

*In vivo*, macrophages are simultaneously exposed to a range of stimuli and have a broad spectrum of functional states, which cannot be divided in simplified M1 and M2 categories, directly corresponding to *in vitro* states. However, even a coarse distinction of *in vivo* macrophages into those with pro-inflammatory and immunosuppressive roles can be of a high clinical value. Hence, we investigated if protein signatures of *in vitro* differentiated M1 and M2 macrophages can be used to study pro- and anti-inflammatory cell populations in patients for which scRNAseq data is available. For this, we obtained publicly available datasets generated for cancer patients within two recent studies that included 15 brain metastases (BrM) samples originating from different primary tumors^16^ and 14 primary hepatocellular carcinoma (HCC) samples^17^. We followed original procedures to separate major cell populations and excluded cells characterized by high expression of tumor markers (EPCAM, KRT19 and MLANA, or AFP, GPC3 and VIL1) from the further analyses (see Methods, as well as Figure 5A and 5B and Figure S4A and S4B). Subsequently, we used the SingleR software tool with Blueprint/ENCODE reference RNAseq profiles from pure cell populations^109,110^ to annotate stromal and immune cells in the tumor microenvironment (TME) (Figure 5A and Figure S4A). Macrophage cells identified this way expressed myeloid markers AIF1, CD14 and LYZ (Figure S4C)^16,17,111^. To further separate macrophages, we composed lists of significantly upregulated proteins in the M1 and M2 *in vitro* states analyzed here. We compared M1 to M2 states and used the top 100 M1 and 100 M2 proteins (Table S6), characterized by high Log_2_FC values and low FDR. The M2 list included proteins significant in both M2a and M2c macrophages. We used the Seurat function ModuleScores for cell annotations (see Methods). Furthermore, we compared these lists to other classification strategies previously applied in the studies of patient macrophages: literature-curated sets of core^112^ and extended^113^ M1- and M2-specific proteins (with 47 and 71 signature entries, respectively, Table S6) and to the CD163 protein alone (at different expression threshold levels). CD163 was used for macrophage classification in recent mass cytometry studies^114,115^.

**Figure 5.**
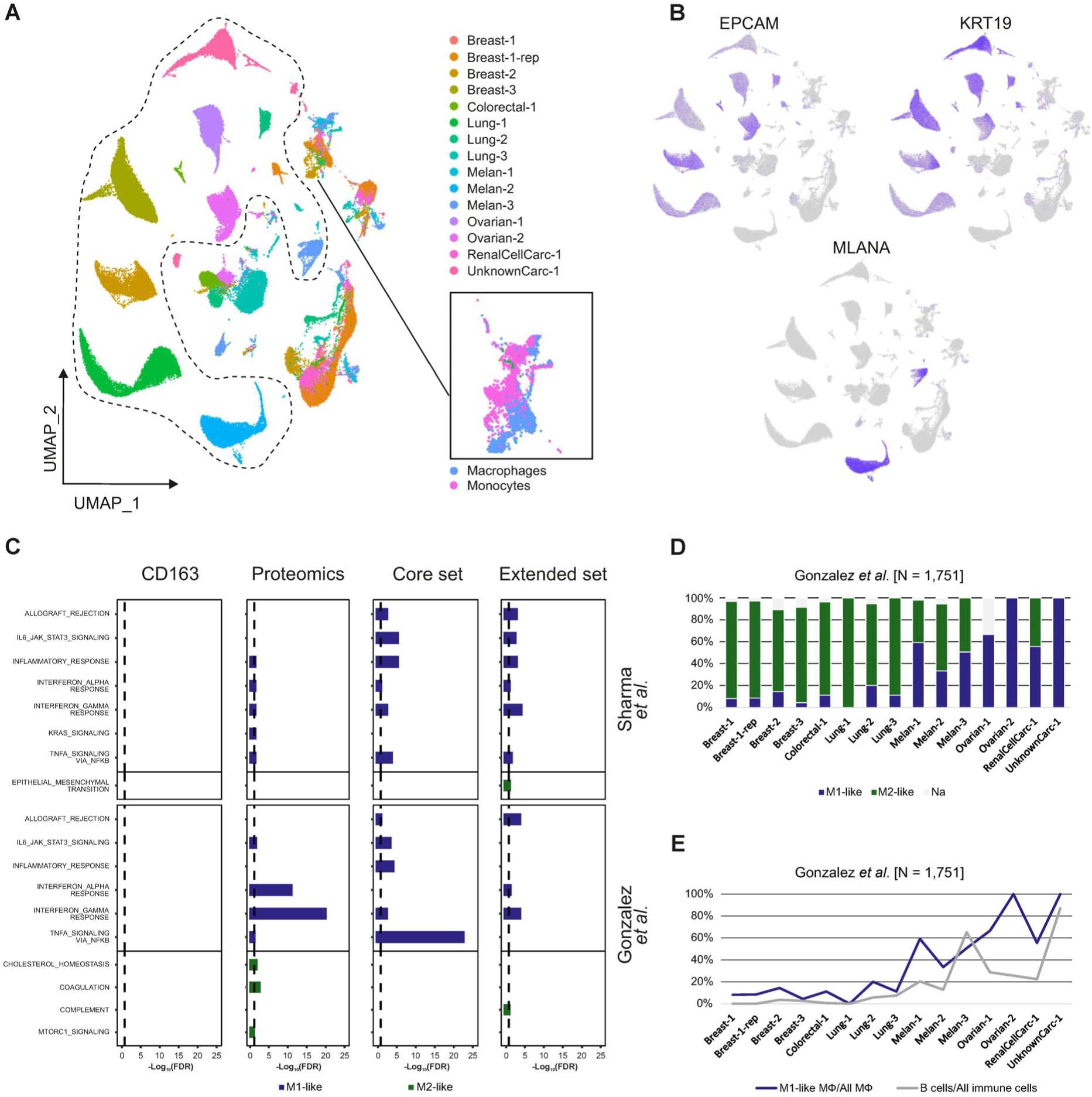
Separation of M1- and M2-like macrophages from clinical scRNAseq data. (A) Two-dimensional visualization of malignant (encircled) and non-malignant single cells in the BrM tumor samples, which were classified based on marker gene expression. Cells from different tumor samples are shown in different colors. A cluster of myeloid cells (see also Figure S4C) is enlarged and macrophages and monocytes, which were assigned according to SingleR annotations, are indicated with different colors. (B) Feature plots of selected malignancy markers, which were used to define tumor cells, are shown. (C) Macrophages were annotated as M1- or M2-like by using different marker signatures (‘Proteomics’ is an unbiased set of markers identified here and the other sets were retrieved from the literatures). For this, scRNAseq data from two studies was retrieved (shown right, Sharma *et al.* corresponds to HCC, Gonzales *et al.* to BrM). MSigDB hallmark terms, which were functionally enriched among the genes upregulated either in M1- or M2-like macrophages, are shown. Significant terms are shown as −Log_10_(FDR) barplots. (D) Percentages of macrophages annotated either as M1-like (blue) or M2-like (green) in each of the BrM sample are shown as a barplot. (E) A fraction of M1-like macrophages (MФ) (among all annotated macrophages) and a fraction of B cells (among all immune cells) are compared across BrM samples. There is a high correlation of the two values (Spearman ***ρ*** = 0.91, p < 2.1×10^−6^).

Depending on the used annotations, we found that up to 89% of the clinical macrophages were classified as M1-like, pro-inflammatory cells (Figure S4D). Furthermore, this analysis showed that the signature proteins defined here through the proteome characterization of *in vitro* macrophages were able to clearly distinguish pro-inflammatory macrophages in both analyzed clinical single cell datasets: Highly expressed genes in the M1-like macrophages, which were classified as such through proteomic signatures, had a strong enrichment in inflammatory functions in both scRNAseq datasets (FDR < 1.49×10^−2^ for the MSigDB hallmark pathway *Interferon gamma response* and *TNFA signaling via NFKB* and FDR < 2.55×10^−2^ for the *Inflammatory response* pathway, Figure 5C, Table S7). Highly expressed genes (with Bonferroni adjusted *p*-values < 0.05, see Methods) were identified through a comparison between the single cells classified as either M1- or M2-like. The inflammatory signal was also strong when we excluded genes used for the classification from the differential expression analysis (Table S7). In addition, macrophages in the scRNAseq-characterized TIME of BrM, which were classified as M2-like through a proteomics-defined signature set, were enriched in the MSigDB hallmark pathway *mTORC1 signaling* (FDR < 3.54×10^−2^). In a comparison, CD163 alone as a marker was not able to classify pro-inflammatory and immunosuppressive macrophages, also when different gene expression thresholds were assessed, but the two knowledge-based lists clearly defined pro-inflammatory clinical macrophage subsets (Figure 5C). This highlights a previous notion that effective macrophage classifications benefit from including multiple markers^112^. Overall, the unbiased list of differentially expressed proteins obtained here had a comparable power in macrophage classification as the literature lists, which were based on evidence from multiple independent studies of macrophage functions (Figure 5C). It was suggested previously that there is a significant scope to expand and refine biomarkers for clinically relevant macrophage populations through multi-omics analyses^112^. Overall, our results suggest that proteomic studies of different *in vitro* states can be one of the instrumental approaches for this.

Furthermore, we investigated if the presence of pro-inflammatory macrophages correlated with the overall TIME composition and the presence of any other cell types. In the BrM tumors that originated from melanoma, which are often more immunogenic but with an exhausted immune signature, we observed a higher fraction of pro-inflammatory macrophages than in samples originating from breast, colorectal and lung tumors (>33% vs ≤20% of all macrophages in the TIME, Figure 5D). The sample size used in the study was too small to statistically evaluate this observation. Furthermore, in the set of BrM samples, we found a strong correlation between the fraction of pro-inflammatory macrophages and B cells (Spearman ***ρ*** = 0.91, p < 2.1×10^−6^, Figure 5E). In HCC samples, the fraction of annotated B cells was overall very low (median of 1.5% across patient samples). B cells have been reported to be important for sustaining melanoma associated inflammation and were proposed as a predictor for survival and response to immune checkpoint blockade therapy^116^. Overall, these analyses underline the value of distinguishing different macrophage subpopulations in clinical TIME analyses in comparison to treating macrophages as a homogenous cell type.

## Discussion

The fine balance between macrophage polarization states is relevant for different metabolic and physiological human processes and its disruption associates with several pathologies^5^. *In vivo* macrophage states are highly complex as these cells not only integrate simultaneous and dynamic exposure to dozens of secreted stimuli, but also signals from cell-cell interactions and mechanical signal transduction. Nonetheless, through simplified *in vitro* models it is possible to map major signaling routes associated with individual clinically relevant stimuli and characterize cells with assays that cannot be used *in vivo*. *In vitro* models have been highly useful for the understanding of macrophage biology in infection, wound healing^117^, autoimmune diseases^118^ and cancer^7^. Even though *in vitro* M2 macrophages are not the same as immunosuppressive TAMs, they do have higher expression of markers often used for pro-tumor TAMs, such as CD163 and CD206. Here we found that they also exhibit higher phosphorylation levels of several kinases and TFs, which were previously associated with macrophages that promote tumor growth, such as MAFB, HSF1, PKACα and PDPK1^66,67,77,119^. Proteomics and phosphoproteomics are becoming widely used for characterizing patient samples and for studying cell signaling pathways^120–123^. Here, we exploited their potential for defining signaling proteins that underlie phenotype changes in primary human macrophages.

Signaling pathways in pro-inflammatory macrophages have been intensely studied and here we could recapitulate well-known regulatory roles of p38, JNK and JAK kinases in the M1 state (Figure 2 and Figure 3A). One of the upstream regulators of these kinases is the RIPK2 kinase^124^, which we observe here to be phosphorylated in its active loop. p38 and JNK kinases promote signaling towards the activation of the NF-κB pathway, while JAK kinases activate STAT TFs^125^. Here, we found that STAT1 TF had a higher phosphorylation level in the M1 state (Figure 2A and 2B, Table S1). In addition, together with RELA and NFKB1, the same TF was predicted as most significant regulator for the genes we found upregulated in the M1 state (Table S4) in the public transcriptome analysis. These kinases and TFs were also predicted as central regulatory nodes of the M1 state in the integrative network analysis (Figure 4A, Table S5)^61^.

Signaling routes in immunosuppressive macrophages are less well described, but of a strong interest because of their clinical relevance^24^. In this vein, recently discovered regulation of immune suppression promoted by RIP1 and PI3Kγ kinases in TAMs has attracted attention for possible therapeutic interventions^22,23^. Here, we found that proteins that play a role in the propagation of the *in vitro* M2 states included PDPK1, PKCα, PKACα, PAK2 and LRRK2 kinases. PDPK1 and PAK2 are known substrates of PKCα, while PKACα and LRRK2 can be phosphorylated by the PDPK1 and PAK2 kinases, respectively (Figure 3D and Figure S3). Some of these kinases were also shown to be able to suppress inflammatory pathways *in vivo*^84,87^, and all of them warrant further exploration of their activity status in a clinical context. For instance, the LRRK2 kinase has been associated with the development of both Parkinson and inflammatory bowel diseases^76^. However, the exact role of this kinase in the disease development is still not clear^126^. Even though highlighted by individual studies, neither PKCα, PKACα, PAK2 nor LRRK2 have been so far defined as central regulators of immunosuppressive macrophage states. In addition, by integrating multi-omics datasets, we identified here FOS, NCOR2 and PPARγ TFs as central regulatory nodes in the M2a interaction network. Literature-based interactomes have a bias for prioritizing better studied proteins which can also affect our observations. Nonetheless, a number of macrophage studies^64,65,99,100,105^ have pointed towards the major roles of these TFs in immunosuppressive macrophages.

Because of their fundamental role in cancer and other diseases, there is a strong interest in finding clinically relevant macrophage populations^127–129^ and identifying markers and regulators of the specific cell states^24,130^. Clinical trials that aim to modulate immunosuppressive macrophages include targeting of the CSF1R receptor, delivery of IRF5 and IRF8 TFs that upregulate pro-inflammatory genes, or activation of toll-like receptors^131–133^. On the example of pro-inflammatory macrophages, we show here that unbiased proteomic signatures can be a powerful means for the categorization of macrophages found in the TIME. Of note, macrophages that promote tumor growth often have phenotypes that do not directly resemble *in vitro* generated M2 macrophages^130,134^.

Design of strategies for rational rewiring of cellular pathways benefits from the mechanistic understanding of signal flow. Here, we focused our analysis on the comparison of signaling activities in pro-inflammatory and immunosuppressive macrophages, as reprogramming between the two states is seen as a highly attractive clinical strategy^13,24^. Multi-omics characterization of primary macrophages in different states together with systematic mapping of signaling cascades in these cells provides a global context of cellular activity, which is of a broad interest in the design of new therapeutics.

## Supplementary Figures

**Figure S1.**
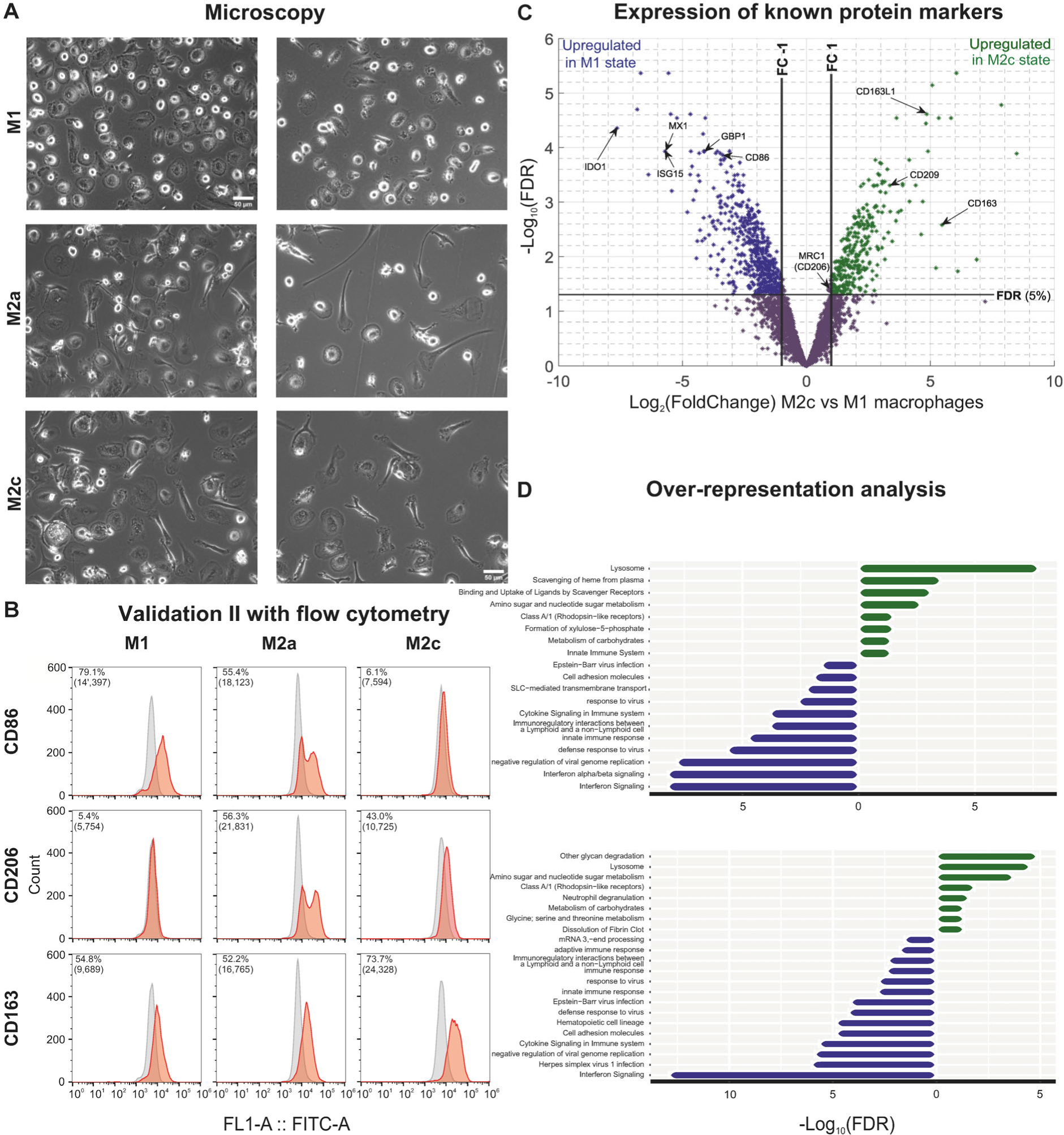
Morphological and molecular properties of primary *in vitro* polarized M1, M2a and M2c phenotypic states. (A) M1, M2a and M2c macrophages were differentiated and polarized from blood-derived monocytes isolated from buffy coats. Shown are images taken after 6 days of differentiation (M1: GM-CSF, M2a/M2c: M-CSF) and 2 days of polarization (M1: LPS/IFN-γ, M2a: IL-4/IL-13, M2c: IL-10). The images were taken from two donors with a Primovert microscope (Carl Zeiss). (B) Flow cytometry profiles for the polarization markers CD86, CD206 and CD163 of *in vitro* polarized macrophages phenotypes in the second of the two tested donors with antigens of interest shown in red and negative or isotype controls shown in grey. Percentages of positive cells, together with the median fluorescence intensity in brackets, are indicated on each individual plot. Y-axes are truncated. (C) Volcano plot illustrating protein marker expressions in terms of Log_2_FC values on the X axis against −Log_10_(FDR) values on the Y axis in a comparison of the M1 and M2c phenotypic states. Differentially expressed proteins are colored blue and green, depending on the directionality of the expression change. (D) Significant (FDR < 0.05) over-representation of differentially expressed proteins (Log_2_FC ≥ 1, FDR < 0.05) between M1 compared to M2a (top) and M2c (bottom) macrophages within KEGG and Reactome pathways. Blue represents significant pathways related to the M1 phenotypic state, while M2a and M2c are represented in green.

**Figure S2.**
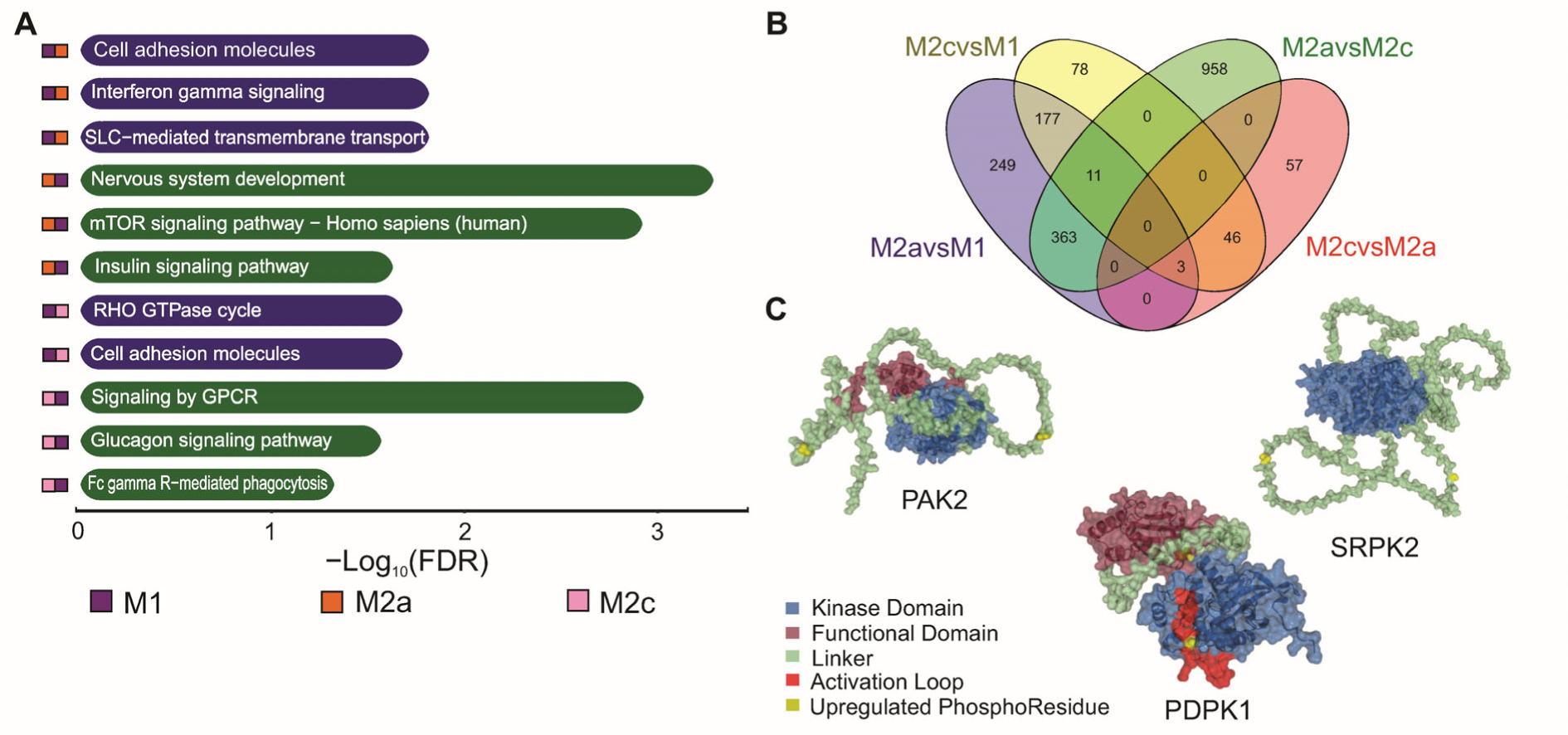
Analysis of phosphoproteins with differential expression levels in the studied conditions. (A) Significant (FDR < 0.05) over-representation of differentially expressed phosphoproteins (Log_2_FC ≥ 1, FDR < 0.05) between M1 compared to M2 macrophages within KEGG and Reactome pathways. Blue represents significant pathways related to the M1 phenotypic state, while M2a and M2c are represented in green. (B) Venn diagram summarizing the differentially expressed phosphoproteins between M2 macrophage phenotypes compared to M1 and between them. (C) 3D protein structure of additional example kinases found important in the current study for immunosuppressive macrophages. The phosphoresidues measured and found upregulated in M2a macrophages are marked in yellow, while different protein functional regions are indicated according to the legend.

**Figure S3.**
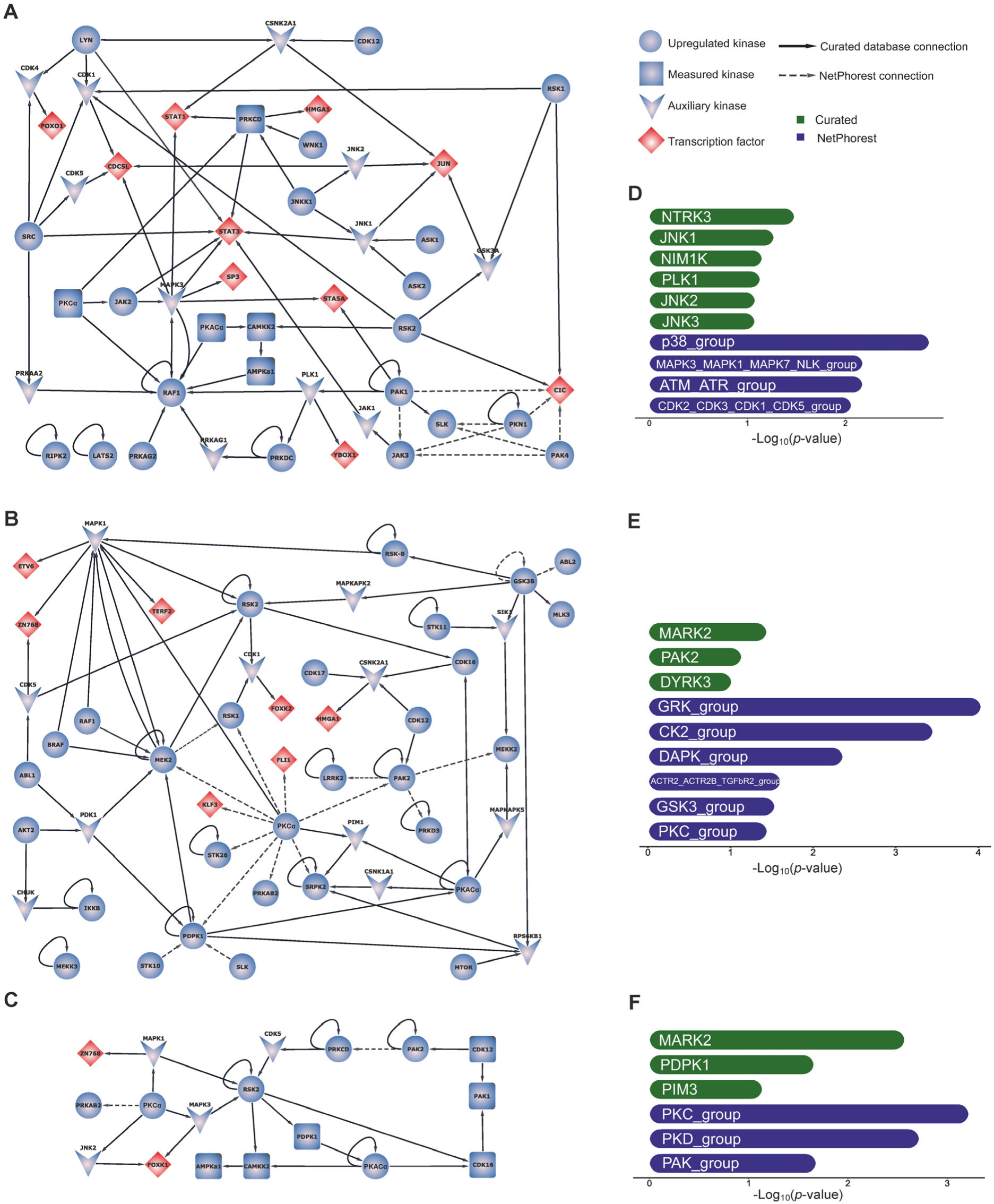
Kinase-Kinase signaling networks and top upstream kinases of each measured macrophage phenotype. (A,B,C) Directional kinase-kinase signaling networks centered around the main upregulated kinases of the M1 compared to M2a (A), M2a compared to M1 (B), and M2c compared to M1 (C) phenotypic state. Within the network, there are modeled kinases with at least one upregulated phosphopeptide (spherical objects) and TFs with upregulated phosphopeptides (rhombic objects). To overcome the challenge of missing values, we allowed the inclusion of known upstream regulatory kinases, which could connect two upregulated kinases or transcription factors in the network, even if they were not measured (V-shaped object) or they were measured but did not have significantly different levels between the states (square objects). Curated kinase-substrate knowledge from PhosphoSitePlus and four other databases was used to connect the kinases (solid lines) as well as the NetPhorest prediction tool to complement the missing knowledge (dotted lines). A connection edge between two upregulated kinases refers to a connection between a kinase that has upregulated phosphorylated residues and a specific peptide that was found to be upregulated as well. This is also valid for the edges with TFs. If an upregulated kinase is linked to a measured but not upregulated kinase, this means that the residues of the latter were not found to be differentially expressed. The presented kinase-kinase signaling maps highlight the signaling transduction routes as were measured in our study. (D,E,F) Upstream kinase activity assessment highlighting the top predicted upstream kinases responsible for the phosphorylation of the upregulated phosphopeptides of M1 compared to M2a (D), M2a compared to M1 (E), and M2c compared to M1 (F) phenotypic state. The analysis was based either on curated phosphorylation databases knowledge or NetPhorest predictions, each analysis relying on a two-sided Fisher’s exact *t*-test where the background was represented by all phosphopeptides measured in the respective phenotypes (see Methods).

**Figure S4.**
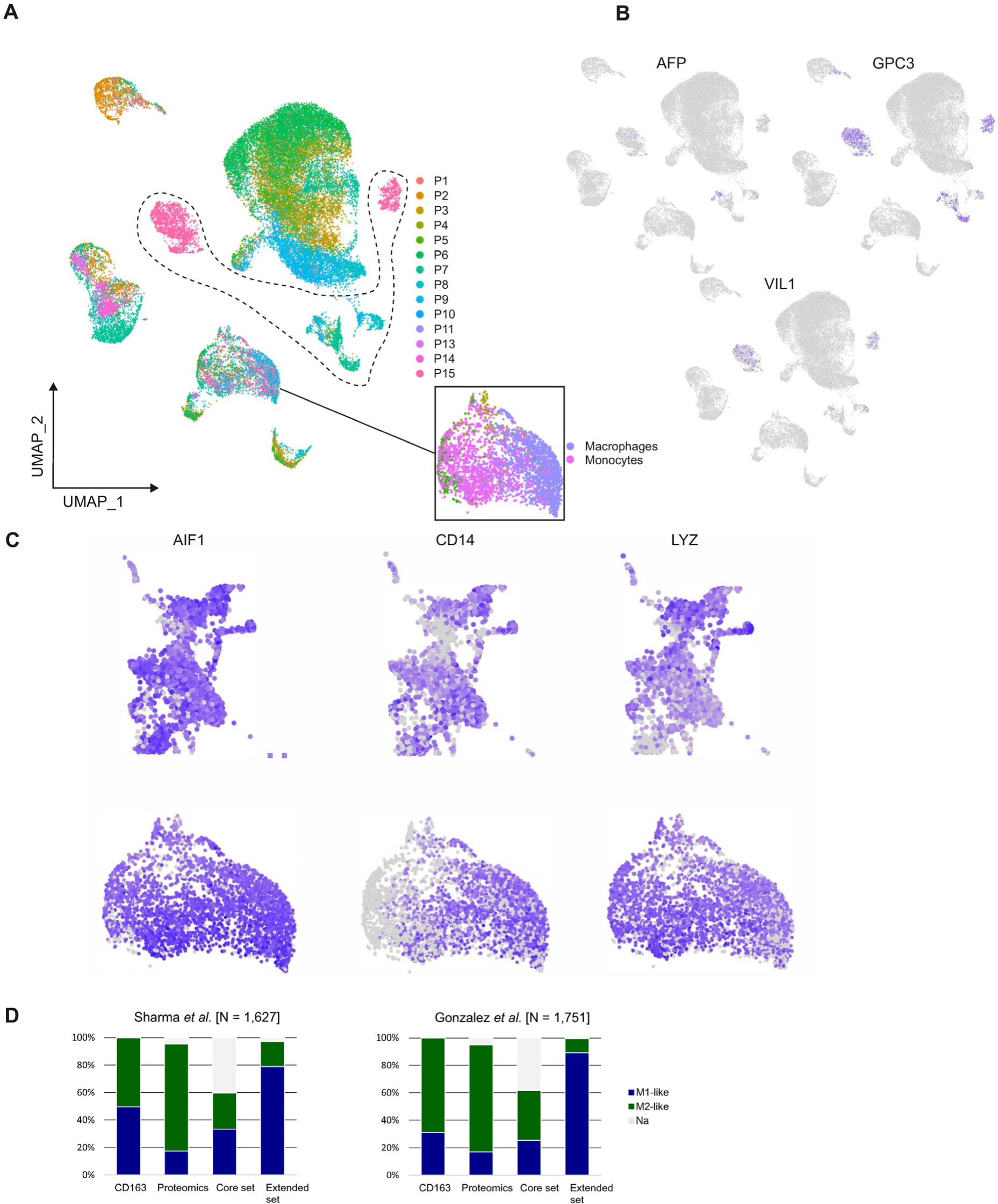
Macrophage annotation using proteomic signatures. (A) Two dimensional visualization of 35’408 malignant (encircled) and non-malignant single cells based on marker gene expression. Colors represent the sample identities. A myeloid cluster (see also Figure S4C) is highlighted with colors representing SingleR annotations. (B) Feature plots of selected malignancy markers. (C) Feature plots of selected myeloid markers of the BrM (top) and HCC (bottom) myeloid cluster. (D) Barplots showing the percentages of annotated macrophages using different marker signatures among the BrM and HCC data sets.

## Methods

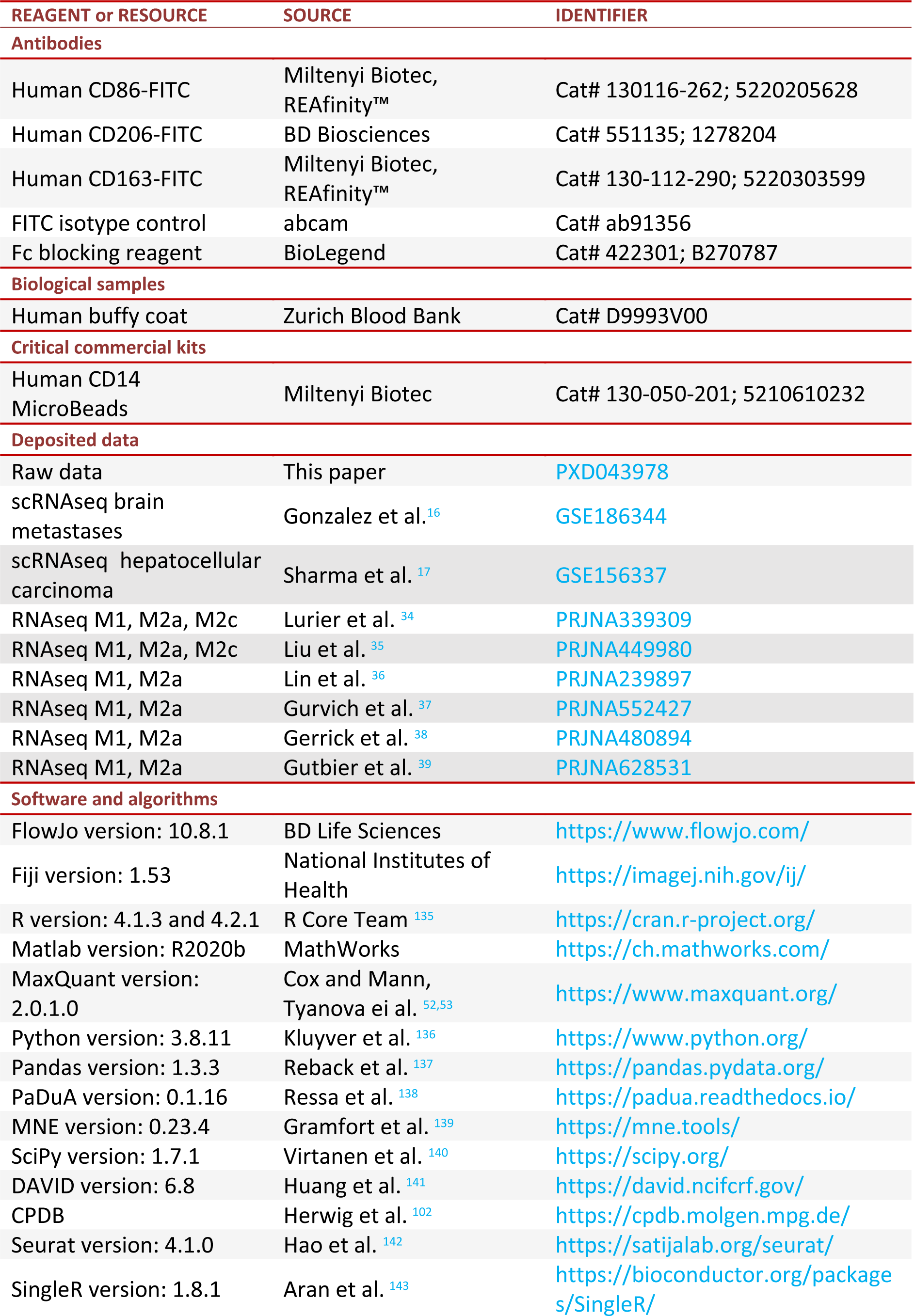
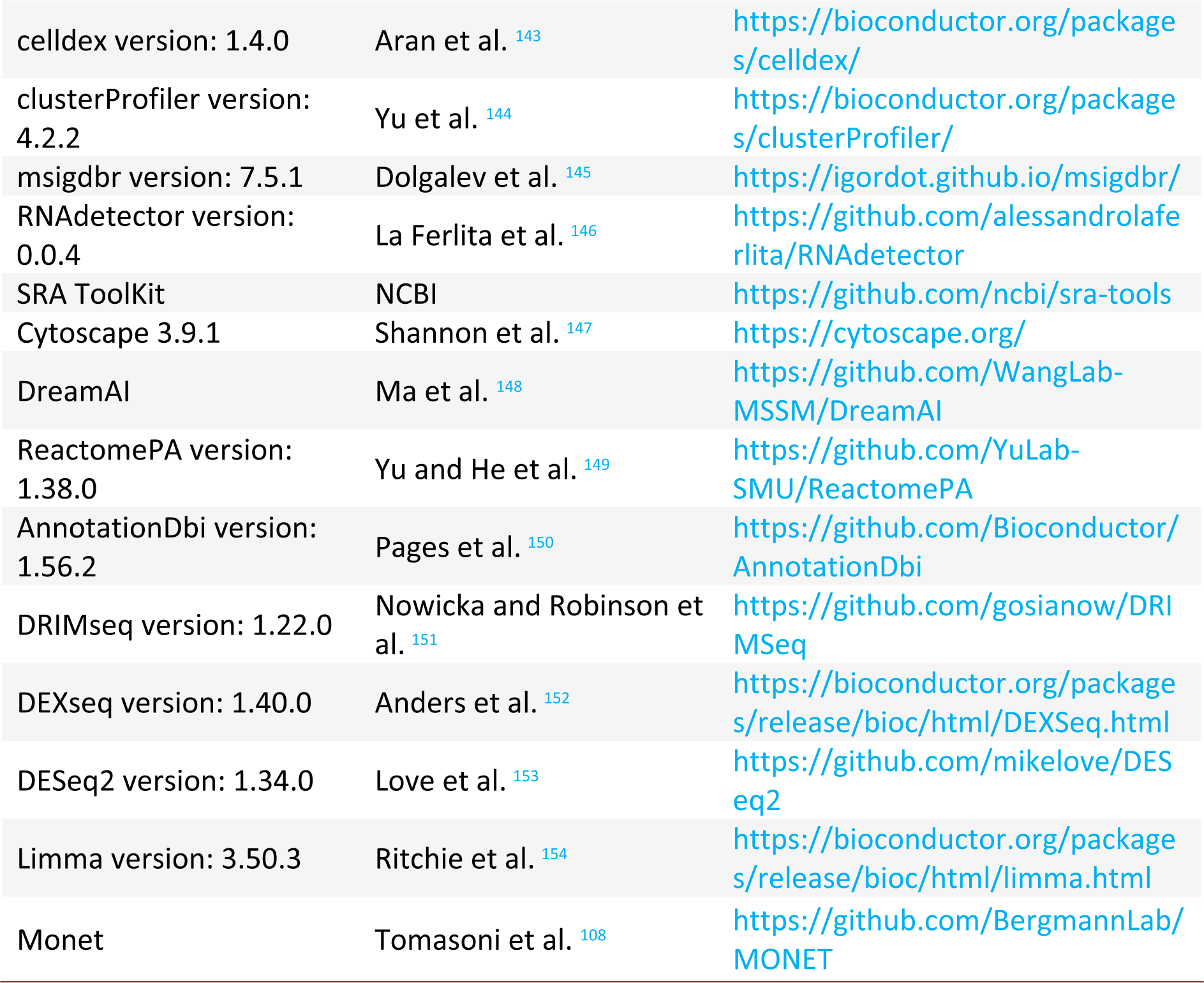

### Experimental workflow

#### Macrophage differentiation and polarization

Buffy coats from four healthy human donors were received after ethics clearance (BASEC Nr. Req_2021-00687) and project approval from the Zurich Blood Bank. Following peripheral blood mononuclear cells (PBMCs) isolation using a density gradient centrifugation with Ficoll (Sigma-Aldrich), monocytes were positively selected with a commercial CD14 kit (Miltenyi Biotec). Following selection, the monocytes were differentiated and polarized into the M1, M2a or M2c macrophage states. For M1 macrophages, the cells were first incubated for 6 days at 37 °C with RPMI-1640 (Sigma-Aldrich) containing 10% fetal calf serum (FCS), 1% penicillin-streptomycin (PS), and 15 ng×mL^−1^ of GM-CSF (Sigma-Aldrich). On day 6, the cell culture media was replaced with fresh media containing 100 ng×mL^−1^ of LPS (Sigma-Aldrich) and 20 ng×mL^−1^ of IFN-γ (Sigma-Aldrich) instead of GM-CSF, and the cells were further incubated for 2 days^8,47^. In the case of M2 macrophages, the monocytes were initially treated for 6 days with 30 ng×mL^−1^ of M-CSF (Gibco), then for 2 days with IL-4 (20 ng×mL^−1^) and IL-13 (20 ng×mL^−1^) (Miltenyi Biotec) for the M2a or 40 ng×mL^−1^ IL-10 (ImmunoTools) for M2c macrophages. Cells were plated in 6-well plates (TPP) at a cell density of 2.5×10^6^ cells/well. After polarization, the cells were washed with DPBS and detached through incubation with a cold harvesting solution (10 mM EDTA (Sigma Aldrich) in DPBS), followed by a mechanical step with a cell scraper (VWR). For each polarization state, four biological replicates were prepared, snap frozen and sent on dry ice to the Functional Genomics Centre Zurich (FGCZ) for proteomic and phosphoproteomic measurements.

### Validation of polarization states

For the validation of phenotypic states, we used the remaining cells from two of the four donors (donor III and donor IV in raw proteome files). Aliquots of circa 5×10^6^ frozen monocytes were thawed, differentiated and polarized as described above. Morphological inspection was conducted by acquiring brightfield images using an inverted light microscope (Zeiss Primovert with Axiocam 105 color) equipped with a 20× objective and Ph1 filter. After harvesting, cells were incubated in a 4% (v/v) paraformaldehyde (PFA) solution for fixation. The PFA solution was discarded after which cells were re-suspended and incubated in DPBS containing 10% FCS. As a second blocking step, 5% Fc blocking reagent (BioLegend) per sample were added. Subsequently, fluorescently labeled antibodies were added. Briefly, CD86 (FITC, Miltenyi Biotec), CD206 (FITC, BD Biosciences) and CD163 (FITC, Miltenyi Biotec) were used. Samples were incubated and afterwards 1.2 mL FACS buffer (0.5 % (w/v) bovine serum albumin (BSA) in DPBS) per 100 µL sample was added. Cell-suspensions were centrifuged, the supernatant was discarded, and cells were re-suspensed in FACS buffer. Measurements were conducted on different flow cytometers. For donor III the CytoFLEX S (Beckman Coulter with CytExpert 2.4, Sheath fluid) was used with default setting without discriminator and neutral density filter. Donor IV was measured on a Gallios flow cytometer (Beckman Coulter with IsoFloq Sheath Fluid) with the following settings; Discriminator FS: 20, Particle size: small, Use of neutral density filter, Read 10,000 cells/sample, FS 100 Gain 1, SS 100 Gain 1, FL1 290 Gain 1, FL6 500 Gain 1, Flow rate: medium, Stop conditions: 10’000 and/or 5 min. Data analysis was performed in FlowJo version 10.8.1, whereby measurements were either compared to isotype controls or negative controls (for REAfinity® antibodies), after gating for live single cells.

### Protein extraction and digestion

Peptide extractions and following measurements for four replicates of macrophages in the M1, M2a and M2c states were conducted at the FGCZ. Cells were lysed adding 100 μL FASP lysis buffer (4% SDS, 100 mM Tris / HCL pH 8.2, 0.1 M DTT) and incubated for 10 min at 95 °C. Next, sonication was performed for 1 min at highest amplitude using high intensity focused ultrasound (HIFU). Sonicated liquids were centrifuged and the protein concentrations were determined using commercial protein assay kits (Thermo Fisher). For each replicate, 600 μg of protein was taken and on-filter digested according to an adapted filter aided sample preparation (FASP) protocol^155^. For this purpose, 200 μL UT buffer (8 M urea in 100 mM Tris/HCL pH 8.2) were added, before the samples were loaded onto ultra-filtration units (Merck, MWCO 30 kDa) and centrifuged at 14’000×g. In a further centrifugation step, the SDS containing buffer was exchanged with 200 μL UT buffer. Reduced thiol groups of cysteine amino acids were alkylated by adding 100 μL iodoacetamide solution (0.05 M in UT buffer) and allowing for 5 min incubation time. Subsequently, the samples were washed five times in total (3×100 μL UT buffer, 2×100 μL TEAB buffer at pH 8). Using 120 μL TEAB buffer containing trypsin (Promega) in a 1:50 (w/w) ratio, proteins were on-filter digested overnight in a wet chamber at room temperature (RT). Afterwards, obtained peptides were eluted by applying centrifugation at 14’000×g for 20 min. For analysis of the proteome, 100 μg of peptides was separated and stored separately, while the remaining volume was almost completely dried (∼5 µL) for enrichment of the phosphopeptides.

### Phosphopeptide enrichment

MagReSyn Ti-IMAC beads (ReSyn Biosciences) were used together with a KingFisher Flex System (Thermo Fisher Scientific) to enrich phosphopeptides^156^. Following the manufacturer’s instructions beads were first conditioned with following washing steps: 2×200 µL of 70% ethanol, 1×100 µL 1 M NH_4_OH and 3×loading buffer (0.1 M glycolic acid in 80% ACN, 5% trilfuoroacetic acid (TFA)). After sample dilution with 200 µL loading buffer, beads, wash solutions and samples were loaded into 96 deep well plates and transferred to the KingFisher system. For phosphopeptide enrichment following steps were carried out: 5 min washing of the beads in loading buffer, 20 min phosphopeptide binding to the beads, 2 min washing in loading buffer, 2 min washing in 80% ACN and 1% TFA, 2 min washing in 80% ACN and 1% TFA, 2 min washing in 10% ACN and 0.2% TFA, followed by 10 min elution of the phosphopeptides from the magnetic beads in 1 M NH_4_OH. Phosphopeptides were dried to completeness and re-solubilized with 10 µL of 3% ACN and 0.1% formic acid for MS analysis.

### Liquid chromatography-mass spectrometry analysis

Analysis was conducted for each sample individually in randomized order. In the experimental setup, the samples were subjected to an Orbitrap Fusion Lumos (Thermo Scientific) equipped with a Digital PicoView source (New Objective). There, the samples were first loaded on a trap column (Waters ACQUITY UPLC M-Class Symmetry C18 Trap Column; 100 Å, 5 μm, 180 μm × 20 mm) which was followed by a second column (Waters ACQUITY UPLC M-Class HSS T3 Column; 100 Å, 1.8 μm, 75 μm × 250 mm). The column temperatures were set to 50 °C. During chromatography, peptides were eluted with a constant flow rate at 300 nL×min^−1^. The following elution scheme, where solvent A was composed of 0.1% formic acid and solvent B of 99.9% acetonitrile in 0.1% formic acid, was applied: The initial gradient of 5% solvent B, which was held for 3 min, was increased within 83 min to a total of 22% solvent B. In the next 10 min solvent B was further increased to 32%. This was followed by a 10 min washing step with increasing solvent B content of up to 95%, which was held for another 10 min. Finally, a re-equilibration step was conducted. After accumulation to an automated gain control (AGC) target value (500’000 proteomics, 400’000 phosphoproteomics), full scan MS spectra (from 300 m/z to 1’500 m/z proteomics, from 375 m/z to 1’500 m/z phosphoproteomics) were acquired in the Orbitrap system, where the resolution was set to 120’000 at 200 m/z and the injection time to a measurement-specific time (40 ms proteomics, 50 ms phosphoproteomics). If a precursor exceeded the intensity of 5’000, it was selected for MS/MS. There, the ions were isolated with a quadrupole mass filter (0.8 m/z isolation window proteomics, 1.2 m/z isolation window phosphoproteomics) and further fragmented by application of higher energy collisional dissociation (HCD) using a normalized collision energy (NCE) of 35. By using an adapted universal method (scan rate; rapid, automatic gain control; 10’000 ions, maximum injection time; 50 ms or 120 ms, charge state screening; enabled, singly unassigned charge states; excluded, charge states higher than seven; excluded, precursor masses previously selected for MS/MS measurements; excluded from selection for 20 s, exclusion window: 10 ppm) fragments were detected in the linear ion trap. Samples were acquired using internal lock mass calibration on m/z 371.1010 Th and 445.1200 Th. Results were collected using the local laboratory information system (LIMS) at FGCZ^157^.

### Proteomics and phosphoproteomics

#### Protein and phosphopeptide identification and label free quantification

The MaxQuant software tool^52,53^ version 2.0.1.0 was used with the enzyme settings set to trypsin and protein identification was performed with the integrated Andromeda search engine^53^. The MS data was searched against a database compiled from Homo sapiens proteome sequences. For this, a UniProt^158^ reference was used (taxonomy 9606, canonical version from 2019-07-09). In the settings, carbamidomethylation of cysteine amino acids was set as a fixed modification, while methionine oxidation and N-terminal protein acetylation were set as variable. Further settings specified the minimal peptide length of 7 amino acids, a maximum of two missed-cleavages and the enzyme specificity of trypsin/P. A maximum FDR threshold was set at 0.01 for peptides and 0.05 for proteins. Label free quantification (LFQ) was enabled, and a 2 min window for match between runs was applied.

### Data cleaning

The proteomics analysis was based on the quantitative matrix of protein intensities produced by the MaxQuant analysis (proteinGroups.txt), and the phosphoproteomics analysis was conducted by using the individual peptide phosphorylation intensities in the MaxQuant output (Phospho_STY_Sites.txt), while only considering the first protein in the protein group to which the phoshopeptide was mapped. In both proteomic and phosphoproteomic analysis, commonly occurring contaminants, such as keratins, trypsin and bovine albumin, as well as the peptides matching the reversed sequences in the decoy database were excluded. In proteome analysis, only proteins that were identified based on two or more measured peptides were kept. In the statistical analyses, only peptides and phosphopeptides with at least two measurements in at least one of the studied states were kept for further analysis. Phosphopeptides of class II which had a localization probability below 0.75 were not considered for the kinase enrichment and related analyses as it was not possible to precisely define the phosphorylated residue.

### Data transformation, centering and imputation

LFQ intensity values in proteomics measurements and raw phosphopeptide intensities were Log_2_ transformed and mean-centered across all samples. Missing values were imputed separately for each macrophage state following concepts of the PhosR method^159^. There, missing values that are consistently absent from all samples in a certain phenotypic state and values missing only in a small fraction of samples in a certain condition are distinguished. In order to impute the missing values, two different normal distributions, both shifted left from the mean of the measured values, were constructed by using the PaDuA library^138^. When ≥50% of the biological replicates in a condition were measured, a distribution with a negative shift from 0.5 of the standard deviation (SD) of the original mean was constructed. When <50% of the biological replicates in a condition were measured, a distribution with a negative shift of 1.8 SD was constructed. In both cases, a width of 0.3 SD was applied. Missing data entries were randomly sampled from either of these distributions depending on the fraction of measured values in the condition. The same approach was implemented for both proteomics and phosphoproteomics data. For phosphoproteomics, the intensities of single, double, triple or higher phosphorylated peptides were analyzed separately. The number of statistical tests was taken into account in the differential expression analysis by considering all performed comparisons in the multiple testing correction. Entries that were differentially expressed in different directions depending on the number of phosphorylated residues in the peptide were removed.

### Differential expression analysis

In order to identify differentially expressed proteins and phosphopeptides, moderated *t*-tests were used for a comparison between the states. The obtained *p*-values were corrected for multiple testing with the Benjamini-Hochberg (BH) method^160^. Entries that had an FDR value below 5% and an absolute Log_2_FC of at least 1 were defined as differentially expressed. In the analysis of the phosphoproteome data, protein abundances of the respective phosphopeptides were further taken into account. The aim was to identify phosphopeptides for which the identified expression level changes between the phenotypes were actually driven by differential protein expression. Following the MSstatsPTM approach^161^, phosphopeptides whose change in abundance between specific conditions was not statistically higher than the abundance change between their corresponding proteins were not considered differently regulated and were removed from further analyses (FDR threshold of 5%). A *t*-test was used in order to compare the abundance change of a specific phosphopeptide (PTM) with the abundance change of its corresponding protein between the same two conditions as in (1):

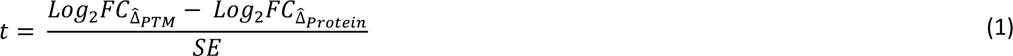

where 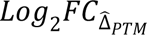 is the binary logarithm of the FC of a specific phosphopeptide calculated from the phosphoproteomics measurements, 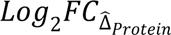 is the binary logarithm of the FC of the corresponding protein calculated from the proteomics measurements, and SE is the standard error of the test statistic described by (2).

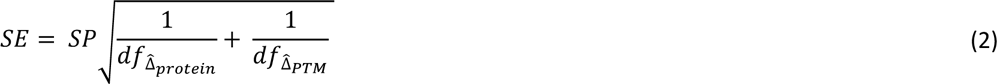

where SP is the pooled standard deviation of the test statistic in accordance to (3), 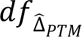 are the degrees of freedom of a *t*-test computed from the phosphoproteomics measurement of the post-translational modification (PTM), while 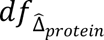 are the degrees of freedom of a *t*-test computed from the proteomics measurement of the corresponding protein.

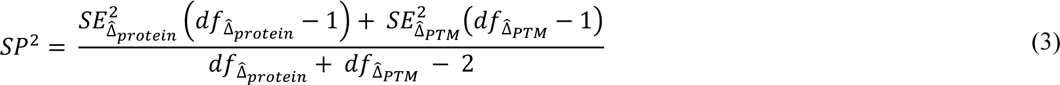

where 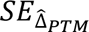 is the standard error of the *t*-test for the differential expression of the respective phosphopeptide between the compared conditions, while 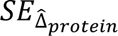 is the standard error of the *t*-test for the differential expression of the corresponding protein for the same conditions.

Finally, the *t*-test statistic was used to distinguish changes in the phosphopeptide level that were not clearly underlined by the changes in protein level. If a protein had at least one phosphopeptide with a change in abundance higher than the change at the protein level, all differentially expressed phosphopeptides of the respective protein were kept for further downstream analysis (i.e. all phosphopeptides found as differentially expressed before comparison to the changes in protein levels).

### Over-representation analysis

In proteins and phosphoproteins that were significantly differentially expressed between macrophage states, over-representation of specific KEGG^162^ and Reactome^163^ signaling pathways was assessed. This was done by using the CPDB^102^ webserver which implements hypergeometric testing for finding statistically significant instances. In addition, the DAVID^141^ tool for functional annotation was used to assess the enrichment in Gene Ontology biological processes. There, a modified Fisher’s exact test is implemented for finding statistically significant instances. In both cases, resulting *p*-values were FDR corrected with the BH method^160^. Background datasets for the comparisons were composed of all measured proteins (for proteome analyses) or included all measured proteins that have annotated phosphorylation residues (for phosphoproteome analyses). For the latter, we used annotations in the PhosphoSitePlus database^97^. Pathways and functional terms identified as being enriched with the FDR threshold of 5% were considered significant. To avoid redundancy, only CPDB significant pathways that had at least two proteins, which were not already assigned to a pathway with a more significant *p*-value, were retained. In order to avoid too specific or non-specific pathways in the phosphoproteome analysis, only pathways that contained at least 5 upregulated phosphoproteins and pathways that did not have more than 300 proteins were retained.

### Phosphoprotein structural conformation

For visualizations shown in Figure 2C and Figure S2C, the full 3D protein structures were extracted from AlphaFold predictions generated with the AlphaFold Monomer v2.0 pipeline^164^. PyMOL version 2.5.2^165^ was used for rendering the protein structures (the script comprising the used parameters is available on the Group’s GitHub page, see below). For each of the presented structures, different structural regions were identified from literature and colored accordingly.

### Inference of kinase activity

Known kinase-substrate relationships with experimental evidence were obtained from PhosphoSitePlus^97^, SIGNOR^166^, PhosphoELM^167^, OmniPath^168^ and PTMSigDB^169^ databases. For the statistical analyses, the number of phosphosites that were found upregulated in each studied state and that could be linked to the upstream kinase were compared to all other phosphosites that were measured in the given phenotype and that could be recognized by the same upstream kinase. Kinases were considered as upregulated when a *p*-value in the two-sided Fisher’s exact test was lower than 0.05. In addition, kinases that could recognize the upregulated phosphopeptides for each state were searched using The Kinase Library, a prediction tool build based on the results from a systematic screen of synthetic peptides^89^. This analysis was performed using the version 0.0.10 of the Kinase Library tool hosted by https://www.phosphosite.org/. We provided the differential expression results for the comparisons between the states with FC and *p*-values calculated as described above. Top ranking upstream kinases in this analysis were identified based on the one-sided Fisher’s exact test with a BH correction for multiple testing. Kinases identified with an FDR < 0.05 were considered as significant. As an alternative method for identifying the most active kinases based on phosphomotifs, we used the NetPhorest prediction tool^95^. NetPhorest predicts kinase families that are able to recognize phosphosites in proteins. For the phosphosites found upregulated here, only predictions with a posterior probability higher than 0.035 as well as those for which the posterior probability was higher than the prior were kept^170^. Only the top three predictions for upstream kinase families (i.e. families with the highest posterior probability) were kept. In order to identify significantly over-represented kinases, we used the same approach as described above and applied a two-sided Fisher’s exact test with multiple testing correction. We considered as significant the kinase groups with 10 or more upregulated substrates and with an adjusted *p*-value below

0.05. Results of these analyses are shown in the Table S3. Finally, the KEA3^96^ algorithm was applied through the online web-service, using as input the upregulated phosphoproteins for each phenotypic comparison. The mean ranks of the KEA3 enrichment scores are listed in the Table S3. All of the above analyses were conducted on the significantly upregulated phosphosites that were identified as differentially regulated with an FDR value < 0.05, for which the observed upregulation was not underlined with the abundance changes in the corresponding proteins and for which the localization probability was higher than 75%. The analyses were conducted in MATLAB R2020b^171^ with the Bioinformatics Toolbox^172^ and Statistics and Machine Learning Toolbox^173^.

### Kinase-Kinase signaling networks generation

A graphical network representation with kinase-kinase and kinase-TF connections was generated using Cytoscape 3.9.1^147^. For this, the information from the curated kinase-kinase interactions and NetPhorest predictions was used. Kinases that were not measured, but that could bridge at least one upregulated kinase and another measured kinase were included in the representation. Additionally, TFs with upregulated phosphosites, which were also known or predicted substrates of the respective kinases were included in the network. Interactions in the network are drawn only when the kinase can recognize the exact phosphopeptides that were found upregulated here. For the larger networks, only upregulated kinases and non-upregulated kinases, which could connect two or more upregulated kinases, were included. The networks were constructed using MATLAB R2020b^171^ together with the Bioinformatics Toolbox^172^ and Statistics and Machine Learning Toolbox^173^.

### RNAseq

#### Data processing and analysis

National Center for Biotechnology Information (NCBI) data repository was searched in order to find transcriptome datasets for primary human macrophages and six bulk RNAseq datasets were downloaded. PRJNA339309^34^ and PRJNA449980^35^ datasets contained transcriptome profiles of all three phenotypes studied here (M1, M2a and M2c, generated with a similar protocol), while PRJNA239897^36^, PRJNA552427^37^, PRJNA480894^38^ and PRJNA628531^39^ datasets contained only measurements for M1 and M2a macrophage phenotypic states. The datasets were pre-processed with the SRA-toolkit 2.11.2 from NCBI, using the fasterq-dump bash commands. The raw FastQ files were then processed using RNAdetector^146^, and aligned to the reference transcriptome of HG38 v33 using the Salmon alignment algorithm^174^. The raw reads were trimmed using Trim Galore^175^, with a minimum read length of 14. The default values were used for the rest of the parameters. Only transcripts with 10 or more counts were considered as expressed and only the protein coding isoforms, based on the annotation in the APPRIS database^176^, were retained.

### Differential gene expression analysis

For each of the obtained transcriptome dataset, differential gene expression analysis was conducted by importing the raw protein coding transcripts’ counts using tximport R package^177^. For M1 and M2a macrophages, only genes whose expression level was in the upper half (i.e. genes that were expressed above the median) in at least 3 different datasets of individual phenotypes were considered. In order to identify significantly differentially expressed genes, DESeq2 R package^153^ was implemented and M1 and M2a macrophages were compared independently in datasets from each of the 6 studies. Entries identified as differentially expressed with a BH adjusted *p*-value < 0.05 and an abundance ratio above four were considered as significant. Following, differentially expressed genes in the 6 analyzed datasets were overlapped and only genes that were differentially expressed in 3 or more datasets were kept. For the M1 and M2c comparison, differentially expressed genes were identified in two datasets using the approach above, and a union of the significant hits in the two studies was used. Following, pathways associated with the identified differentially expressed genes were assessed by using the REACTOME and KEGG database annotations. The background of a specific phenotype was composed from all genes measured in the respective phenotype and pathways identified as significantly over-represented with an FDR threshold <0.05 were considered significant. To avoid redundancy, only CPDB significant pathways that had at least two proteins, which were not already assigned to a pathway with a more significant *p*-value, were retained. Similarly, transcription factors whose downstream targets were over-represented in the sets of differentially expressed genes were identified using TRRUST annotations^101^ available from the CPDB database webtool.

### Differential transcript usage analysis

For differential transcript usage (DTU analysis), raw counts for each dataset were imported with the tximport R package^177^ using the scaledTPM option. Only the transcripts with: (i) higher than a median value in at least 3 datasets, (ii) a minimal proportion of 0.05, and (iii) the corresponding gene expressed in all replicates in the original studies, were considered. The DTU analysis was conducted with the DRIMseq R package^151^ using the add_uniform parameter. In order to extract genes that contained evidence of DTU between the assessed macrophage phenotypes, a two-stage correction method in the StageR R package with an FDR threshold of 0.05 was applied^178^. Following, genes that had the DTU evidence were overlapped between all the analyzed datasets and only the ones that were found in at least half of the comparisons were retained. Next, the enriched pathways were identified analogously to the differential gene expression analysis.

### Protein-Protein interaction networks

We used publicly available interaction data in order to investigate connections among the entries identified as significantly differentially expressed in the phosphoproteomics, proteomics, transcriptomics and alternative splicing (i.e. DTU) analysis. For this, we obtained high confidence human interactors from the STRING^179^, BioGRID^180^ and IntACT^181^ databases. For STRING, database version 11.5 containing the complete interactions data considering all sources was used and filtered to include only entries with a combined score above 0.7. For the BioGRID version 4.4.218, data file in the mitab format that contained a dataset of interactors with physical interactions supported by independent validations was used. The latter file was filtered to exclude entries without an associated confidence value. For the IntACT database, the psimitab from 13/07/2022 was used and filtered to keep only interactions with a confidence score above 0.7. Interaction pairs obtained from the different databases were overlapped and merged in a joint dataset.

The obtained interaction data was used to construct a network. For this, only interaction pairs in which both of the entries were found among significantly differentially expressed hits in at least one of the analyses (transcriptomics, DTU, proteomics or phosphoproteomics) were kept. Genes with significant DTU did not show as strong enrichment in the expected M1/M2 processes as other hits, so networks composed exclusively from the DTU hits (>75% entries) were discarded. The network with the highest number of connected members was analyzed further.

In order to identify central nodes in the network, which are able to most effectively connect other network elements, current flow betweenness centrality metric was used^104^. The networks were analyzed using the centiserve^182^, CINNA^183^, igraph^184^ and tidygraph^185^ R packages.

Network modules with more closely connected entries^186^ were extracted from the analyzed network using the MONET software^108^. For this, the Modularity optimization method with undirected edges was used, and the desired average nodes degree in the identified modules was set to 10. The modules were sorted based on the number of nodes they included. Next, Reactome and KEGG pathways over-represented in individual modules were assessed. Background datasets for individual phenotypes were composed of all genes and proteins that were used to construct the full-scale network of the respective phenotypes. Pathways with at least five significant hits and an FDR < 0.05 were considered significant. Less significant redundant pathways as well as non-specific pathways with more than 300 members were omitted from the final report (following the approach described above).

The clusterProfiler^187^ R package was used for the pathway analysis. For the visualization of results, circular charts were generated with the circlize R package^188^. For this, only the top 30% most central nodes (based on the current-flow betweenness centrality) were represented. For data handling and visualization, additional R packages were used: RColorBrewer^189^, readr^190^, stringr^191^, gtools^192^, gridBase^193^, ComplexHeatmap^194^, tidygraph^195^, biomaRt^196^ and readxl^197^.

### scRNAseq

#### Data processing

TME scRNAseq datasets were downloaded from the previously published studies deposited at the NCBI Gene Expression Omnibus (GEO). The datasets corresponded to TME of 14 primary HCC samples (accession number GSE156337)^17^ and 15 BrM^16^. The latter metastases originated from different primary tumors (accession number GSE186344). Data was imported into R^135^ and handled using the Seurat package^142^. Outliers were excluded from unfiltered datasets by removing cells with less than 500 or more than 9,000 expressed genes and cells with more than 10% mitochondrial genes. Furthermore, genes expressed in fewer than three cells were not used for the subsequent analyses. Feature counts (i.e. counts per gene) were normalized through division by the total counts of the corresponding cell and were then multiplied by a scale factor of 10,000. Following, the normalized counts were natural-log transformed. The 2,000 most variable genes were identified by variance stabilizing transformation. Next, percentage of mitochondrial genes and sequencing depth, which can artificially drive cell clustering, were regressed out against each feature (i.e. gene) using the function *vars.to.regress*. Subsequently, residuals were scaled and centered to a mean expression of zero and a variance of one across cells.

### Separation of stromal and immune cells

Based on the identified 2,000 most variable genes, principal component analyses (PCA) were conducted for each dataset individually, where the dimensions were first reduced to 40 principal components (PC). In order to determine suitable numbers of PCs for further dimensionality reductions, elbowplots were generated using the uniform manifold approximation and projection (UMAP) algorithm. It was decided to proceed with 13 and 18 PCs for the HCC and BrM datasets, respectively. For clustering, shared nearest neighbor (SNN) graphs were constructed based on the euclidean distances in the PCA space. Cells were grouped together with the Louvain algorithm, with resolutions between 0.1 and 1.5, whereas appropriate values were selected individually based on visual inspection. Feature plots of the selected markers were used to separate and extract stromal and immune cells from the remaining highly variable cancer cells. The markers originally used by Gonzalez *et al*. were implemented here for the BrM dataset (T cells: CD3D, IL7R; B cells: JCHAIN, MZB1; Endothelial cells: CLDN5, PECAM1; Astrocytes: GFAP, S100B; Dendritic cells: CD1C, CLEC10A; Macrophage: AIF2, LYZ; Mesenchymal cells: ISLR, CTHRC1; Mural cells: RGS5, ACAT2; Cancer cells: MLANA, KRT19, EPCAM), whereas the markers used in the original study by Sharma *et al*. were also implemented here for the analysis of the HCC dataset (T cells: CD3E, IL7R; B cells: MZB1, CD79A; Endothelial cells: PECAM1, VWF; Fibroblasts: ACTA2, THY1; Hepatocytes: ALB, KRT8; Myeloid cells: LYZ, CD14; NK cells: GNYL, NKG7; Cancer cells: AFP, VIL1, GPC3).

### Extraction of macrophages

Extracted stromal and immune cells were further annotated using the SingleR package^143^. The annotation was based on an altered version of the celldex^143^ reference dataset Blueprint/ENCODE^109,110^. The aim was to retain only cell types expected in the specific tissue and in this way reduce the noise. Cells, which were annotated as macrophages based on the reference bulk RNAseq data from pure cell populations, were extracted for the further analysis.

### Macrophage annotation

In the HCC and BrM datasets, each macrophage was individually assigned to either the M1- or M2-like polarization status according to four different criteria. First, expression of the CD163 receptor gene was used as a criteria to designate macrophages to the M2- or M1-like category, depending if the gene was expressed or not, respectively. In addition to this base analysis, different thresholds for the CD163 expression levels were used for the M1- and M2-like categorization. Second, signature sets for the cell classification were constructed based on the top 100 most highly upregulated proteins in the M1- and M2-like polarization states. The proteins were identified from the comparison of the macrophage proteomics profiles and were all significantly differentially expressed between the states (FDR < 0.05) (Table S6). Third, a signature set of M1- and M2-like markers, which represented a consensus in the community was retrieved from literature^112^ (referred to as core set). Fourth, a set of M1- and M2-like markers, which was collected from the literature for the previous study^113^ (referred to as extended set) was used. All signature sets are listed in Table S6. Each single cell was classified according to a ModuleScore, which was calculated for signature proteins. This was repeated independently for three different signature sets. For each set, ModuleScores were calculated based on the comparison of gene expression levels of the signature genes on one side and a control set of genes with a similar average expression across all cells on the other side. To identify the latter set, genes in the single cells classified as macrophages were binned according to their average expression levels across all cells in 24 bins. For each gene in the signature list, 100 control genes were randomly selected from the same expression bin. Next, on single cell level, from the expression value of each gene, the average expression values of the corresponding randomly selected control genes were subtracted. The average expression value of all genes in the signature list was then calculated for each cell, which yielded the gene set activity estimate for a single cell. For each reference set analysis, cells were assigned to the M1- or M2-like polarization state when the gene set activity was higher in one of the states and when it had a positive value. When both of the gene set activity scores were negative, cells were categorized as an unknown (*Na*) macrophage group. Each macrophage was independently annotated as either M1-, M2-like or *Na* according to four different criteria listed here.

### Differential expression analysis of scRNA datasets

In order to identify differentially expressed genes between the here-annotated M1- and M2-like macrophage groups, a differential expression analysis was conducted. Differentially expressed genes were identified based on student *t*-tests followed by a Bonferroni correction for multiple testing. Differentially expressed genes with an adjusted *p-*value < 0.05 for a comparison between the states and an average Log_2_FC > 0.75 were considered significant. For the latter analysis, only genes which were expressed in at least 10% of cells in either of the states were assessed. Lists of differentially expressed genes (Table S6) were then analyzed with the clusterProfiler^144^ and msigdbr^145^ packages in order to identify over-representation of functionally related gene sets annotated within the hallmark collection (Table S7). There, a BH corrected *p*-value <0.05 and a gene count above one were used to define significant terms.

## Data and code availability

Proteomics and Phosphoproteomics data are available within the PRIDE repository^198^ under the accession number: PXD043978 (Username, Password: Bg60lqYr). Code used in this study is available on the GitHub code repository https://github.ch/MOFHM/MacrophageSignaling.

## Acknowledgments

The authors gratefully acknowledge the Functional Genomics Center Zurich (FGCZ) of University of Zurich and ETH Zurich, and in particular Dr. Antje Dittmann and Laura Kunz, for the support on proteomics and phosphoproteomics analyses. Figure 1A and the graphical abstract were created with BioRender.com.

## Author contributions

T.T. performed data analysis, results interpretation and visualization for the phosphoproteomics, public transcriptomics and network integration part. J.B. performed data analysis, results interpretation and visualisation for the proteomics and public scRNAseq part and he performed *in vitro* macrophage polarization and flow cytometry experiments. V.A-N. performed monocyte isolation, *in vitro* macrophage polarization and flow cytometry experiments. V.A-N., K.H, B.S., G.Y, C.L and M.R. contributed to the experimental design, troubleshooting, interpretation of results and text writing. M.B. conceptualized and supervised the study. M.B, T.T and J.B. wrote the manuscript with the input from all authors.

## Supplementary tables

Table S1: Differential expression analysis of proteomics and phosphoproteomics data

Table S2: Pathway over-representation analysis for the differentially expressed proteins and phosphoproteins

Table S3: Kinase enrichment analysis

Table S4: Transcriptomics analysis of six RNAseq datasets

Table S5: Network analysis of integrated multi-omics datasets (Transcriptomics + Phosphoproteomics + Proteomics)

Table S6: Differentially expressed genes between M1- and M2-like macrophages

Table S7: Over-representation analysis of HALLMARK signatures within differentially expressed genes between M1- and M2-like macrophages

